# Engineering of CRISPR-Cas PAM recognition using deep learning of vast evolutionary data

**DOI:** 10.1101/2025.01.06.631536

**Authors:** Stephen Nayfach, Aadyot Bhatnagar, Andrey Novichkov, Gabriella O. Estevam, Nahye Kim, Emily Hill, Jeffrey A. Ruffolo, Rachel Silverstein, Joseph Gallagher, Benjamin Kleinstiver, Alexander J. Meeske, Peter Cameron, Ali Madani

## Abstract

CRISPR-Cas enzymes must recognize a protospacer-adjacent motif (PAM) to edit a genomic site, significantly limiting the range of targetable sequences in a genome. Machine learning-based protein engineering provides a powerful solution to efficiently generate Cas protein variants tailored to recognize specific PAMs. Here, we present Protein2PAM, an evolution-informed deep learning model trained on a dataset of over 45,000 CRISPR-Cas PAMs. Protein2PAM rapidly and accurately predicts PAM specificity directly from Cas proteins across Type I, II, and V CRISPR-Cas systems. Using *in silico* deep mutational scanning, we demonstrate that the model can identify residues critical for PAM recognition in Cas9 without utilizing structural information. As a proof of concept for protein engineering, we employ Protein2PAM to computationally evolve Nme1Cas9, generating variants with broadened PAM recognition and up to a 50-fold increase in PAM cleavage rates compared to the wild-type under *in vitro* conditions. This work represents the first successful application of machine learning to achieve customization of Cas enzymes for alternate PAM recognition, paving the way for personalized genome editing.

## Introduction

The protospacer-adjacent motif (PAM) is a short DNA sequence next to a target site that CRISPR-Cas proteins must recognize to bind and cleave DNA. PAM binding is essential for initiating DNA unwinding, R-loop formation, and efficiently locating a genomic target (1). In nature, bacteria-phage co-evolution has driven the diversification of Cas proteins, enabling them to recognize a wide range of PAMs (2–4). In genome editing, the PAM is essential for specificity but restricts the range of genomic sites available for editing. This poses challenges for modalities like base editing and homology-directed repair, where precise positioning of the Cas protein is critical (5). In contrast, strict PAM recognition can be leveraged for applications requiring high specificity, such as single-nucleotide allele discrimination and the precise targeting of dominant-negative disease-associated mutations (6).

A variety of experimental approaches have been developed to engineer CRISPR-Cas enzymes with altered PAM specificity. Rational engineering approaches have focused on mutating key PAM-interacting residues to broaden (7–10) or shift PAM recognition (11). For example, structure-guided mutagenesis enabled the engineering of near-PAMless CRISPR-Cas9 enzymes capable of editing most sites in the human genome (8). Experimental evolution methods – such as phage-assisted continuous evolution (PACE) (12–14) and bacteria-based selection (10) – have also been employed to broaden PAM specificity but require labor-intensive and iterative experimentation. Despite these advances, there is still a need for a robust and facile method to engineer Cas enzymes with customized PAMs for specific therapeutic targets and scalable personalized medicine.

Large language models provide a powerful framework for protein engineering (15, 16), including for genome editors (17, 18). In this study, we explored their potential to predict and customize PAM recognition for CRISPR-Cas proteins. To achieve this, we compiled a large and diverse training dataset of CRISPR systems and their associated PAMs through systematic genome data mining (17). Using this dataset, we developed Protein2PAM, a machine learning framework that can accurately predict PAMs directly from diverse Cas protein sequences. We demonstrate that Protein2PAM had learned biophysical principles of PAM recognition and can identify PAM-interacting residues in Cas proteins without utilizing structural information. Additionally, we show that Protein2PAM can be utilized to generate highly active, PAM-customized enzyme variants in a single step, without iterative experimentation. To support PAM identification and accelerate genome editing for the scientific community, we have made Protein2PAM freely available at https://protein2pam.profluent.bio.

## Results

### Evolutionary landscape of CRISPR-Cas PAMs

No comprehensive dataset for CRISPR-Cas PAMs existed at the time of this study, limiting the ability to model how Cas proteins interact with their PAMs. To overcome this, we conducted extensive data mining of 26.2 Tbp of assembled microbial genomes and metagenomes to build the CRISPR-Cas Atlas (Fig. 1a) (17). We identified PAMs for CRISPR-Cas Types I, II, and V, which are DNA-targeting systems that utilize a PAM during target interference and were well represented in the CRISPR-Cas Atlas. We did not predict PAMs for CRISPR-Cas Types III and VI, which target RNA and avoid self-immunity through PAM-independent mechanisms (19, 20), as well as Type IV, which was poorly represented in the CRISPR-Cas Atlas. To identify PAMs, we searched for the natural targets of CRISPR spacers in a database of over 16 million virus and plasmid genomes (21, 22) and looked for conserved motifs flanking protospacers (23). This process resulted in 45,816 distinct PAM predictions which formed our training dataset and covered 71.6% of CRISPR-Cas operons (Fig. 1b).

**Figure 1.**
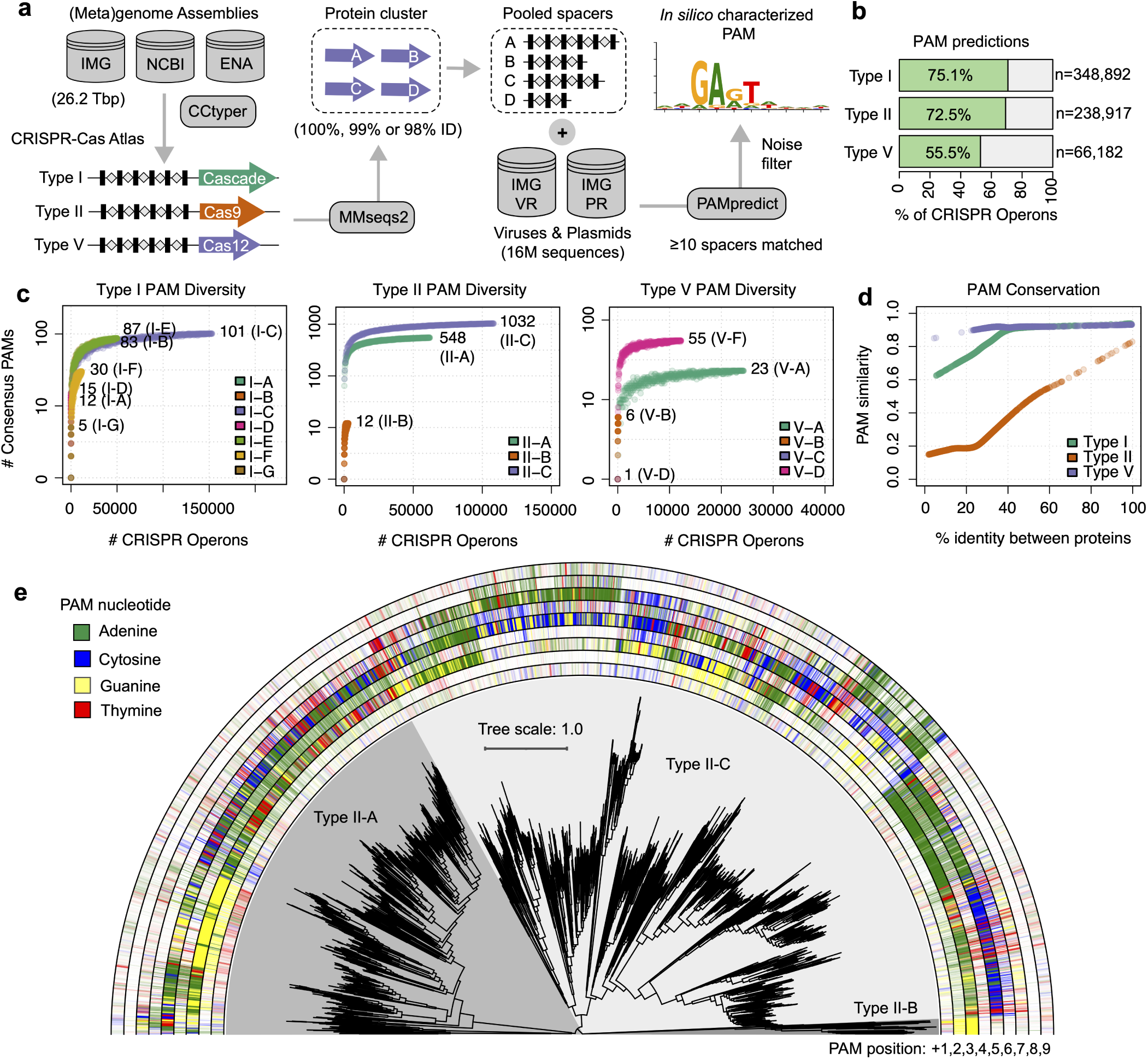
Systematic identification of CRISPR-Cas PAMs. (a) A bioinformatics pipeline was employed to identify PAMs across diverse CRISPR-Cas systems. The pipeline aligned CRISPR spacers to a large database of viral and plasmid sequences to detect conserved flanking motifs. The Cas proteins responsible for PAM recognition during target inference are shown: Cas9 and Cas12 function as single-protein effectors, while Cas8 operates as part of the multi-subunit Cascade complex. In total, 45,816 distinct PAM predictions were made (Type I: *n* = 28, 410, Type II: *n* = 15, 731, Type V: *n* = 1, 675). (b) Fraction of CRISPR-Cas operons associated with a PAM prediction. (c) Accumulation curves of PAM diversity with increasing data volume. Discovery of new PAMs has largely plateaued for Type I and II systems. (d) PAM similarity was compared between Cas proteins with different levels of relatedness. PAM similarity rapidly diverges for Type II systems but is highly conserved for Types I and V. (e) A phylogenetic tree of Cas9 proteins clustered at 70% identity using MMseqs2 (25). Outer rings indicate the information content at each of the first 9 PAM positions. Phylogenetic tree built using FastTree (26) and visualized using iToL (27).

Our dataset represents a 2.8-fold increase over the largest dataset of bioinformatically determined Cas9 PAMs (23) and a ∼200-fold increase over the largest dataset of experimentally determined Cas9 PAMs (3) (Methods). Collectively, the PAMs in our dataset have the potential to cover all possible 10-bp regions, with each site being targetable by a median of 648 Cas enzymes in our training dataset. Discovery of new PAMs has plateaued for most CRISPR subtypes, suggesting that our dataset captures a majority of PAM diversity in nature (Fig. 1c). The exception was Type V systems, which had the lowest PAM prediction rate (Fig. 1b) and where PAM predictions were determined for only four of fifteen literature reported subtypes (24).

Type II CRISPR systems displayed the highest PAM diversity, representing 81.6% of unique consensus PAMs. Further, Type II PAMs evolved rapidly over short evolutionary distances (Fig. 1d-e), whereas PAMs for Type I and Type V systems were highly conserved (Fig. 1d and Fig. S1). While not fully understood, this difference in PAM variability likely reflects distinct evolutionary pressures on each CRISPR-Cas system and enables Cas9 to more rapidly adapt to evolving threats from phages and mobile genetic elements.

### A machine learning framework to predict PAMs from Cas proteins

Next, we leveraged protein language models (pLMs) to learn the relationship between Cas proteins and their PAMs (Fig. 2a-b). For each CRISPR-Cas type, we selected the protein family responsible for PAM recognition during target interference: Cas8 for Type I (or Cas10d for Type I-D), Cas9 for Type II, and Cas12 for Type V (28). Cas9 and Cas12 function as single-protein effectors, while Cas8 operates as part of the multi-subunit Cascade complex. The PAM was represented as a sequence of 10 probability vectors over the nucleotides A, C, G, and T, located either upstream (Types I and V) or downstream (Type II) of the protospacer. Our approach assumes that the nucleotides in the PAM are conditionally independent, given the protein sequence, which is supported by experimental evidence that specific residues in the protein interact with individual nucleotides within the PAM (28).

**Figure 2.**
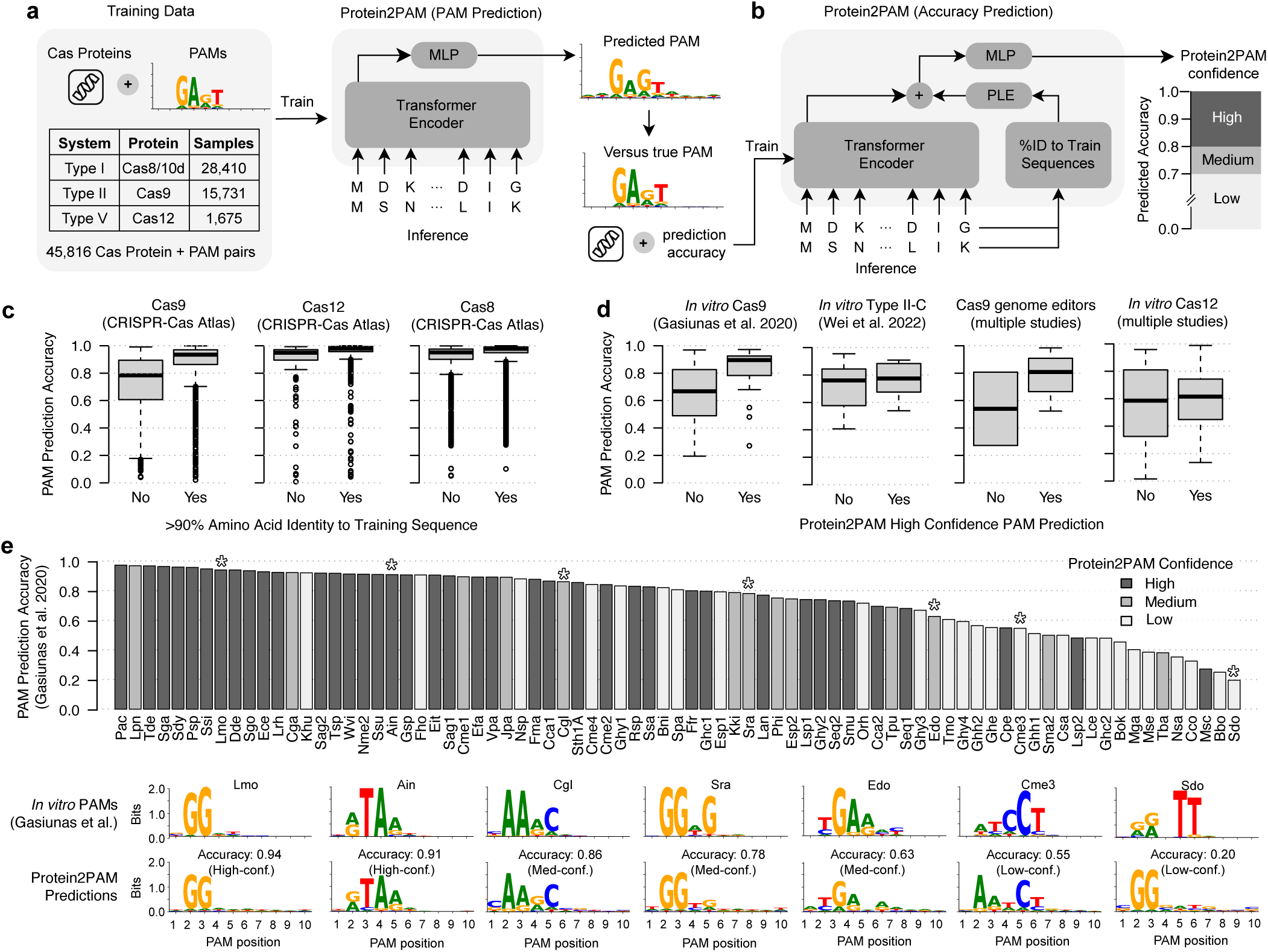
A machine learning framework to predict PAMs from Cas proteins. (a) The Protein2PAM model architecture consists of a pre-trained 650-million-parameter transformer encoder (16), followed by a 2-layer multi-layer perceptron (MLP) head responsible for predicting PAM nucleotide probabilities. (b) Architecture of model for quantifying Protein2PAM’s confidence in its own predictions, incorporating both protein language model (pLM) embeddings and distance to training sequences. (c) PAM prediction accuracy for protein:PAM pairs held back from the CRISPR-Cas Atlas training dataset. (d) Prediction accuracy for Cas proteins with experimentally characterized PAMs. (e) PAM prediction accuracy for 79 diverse Cas9 orthologs experimentally characterized in Gasiunas et al. (21). Representative examples are indicated below the barplot. In all panels, PAM prediction accuracy was measured using cosine similarity.

The Protein2PAM model architecture consisted of a pre-trained 650-million-parameter transformer encoder (16), followed by a 2-layer multi-layer perceptron (MLP) head responsible for predicting PAM nucleotide probabilities (Fig. 2a). The transformer captured key dependencies between amino acid residues relevant for PAM recognition, and the [*CLS*] token from the encoder’s final layer was passed to the MLP, which output a 10 x 4 matrix representing the predicted nucleotide probabilities at each PAM position. Protein2PAM models for Type I and V rapidly converged to their minimum loss, while the more variable PAM recognition in Type II systems led to longer training times and a higher final loss (Fig. S2). In addition to the PAM prediction model, we trained a separate model that estimates PAM prediction accuracy, incorporating both pLM embeddings and amino acid identity to training sequences (Fig. 2b and Fig. S3).

Next, we investigated the optimal input sequence for modeling (Fig. S3a). In Type II systems, PAM recognition is primarily mediated by Cas9’s PAM-interacting domain (PID) (28, 29). We used a custom Hidden Markov Model (HMM) database to identify PID regions and trained a separate model on these sequences. The PID-only model outperformed the full-sequence Cas9 model, likely due to more effective feature selection. In Type I systems, evidence suggests that Cas5 may also contribute to PAM recognition (28). However, including Cas5 alongside Cas8/10d in the model reduced accuracy, especially for sequences more distant from the training data. Based on these results, we selected the Cas8/10d-only model for Type I, the PID-only model for Type II, and the full-sequence model for Type V, as these configurations demonstrated the best generalization to new data.

Protein2PAM neural models demonstrated high accuracy in predicting PAMs for diverse CRISPR-Cas systems, with accuracies of 0.949 for Type I, 0.868 for Type II, and 0.955 for Type V systems (Fig. 2c). Accuracy was measured using the cosine similarity between PAMs predicted by the model and PAMs held out from the CRISPR-Cas Atlas training dataset (Methods). Protein2PAM models were considerably more accurate than a baseline method that predicted PAMs based on the PAM of the nearest protein sequence in the training set (Fig. S3a). For proteins with less than 90% sequence identity to the training data, the Type II model showed a drop in accuracy, while the Type I and V models remained relatively stable (Fig. 2c), reflecting the dynamics observed during model training.

### Model concordance with *in vitro* determined PAMs

To more robustly evaluate Protein2PAM, we applied the models to Cas proteins with experimentally determined PAMs (Fig. 2d-e and Table S1). We first applied Protein2PAM to 14 diverse Type I CRISPR and CAST systems experimentally characterized by Wimmer et al. (30). Using Cas8 proteins as input, Protein2PAM successfully recapitulated consensus PAMs for every active CRISPR system in the study (Fig. S4), including for proteins with as low as 25% amino acid identity to a training sequence.

Next, we applied Protein2PAM to 112 Type II systems, predicting PAMs for diverse Cas9s (3), closely related Cas9s (31), and Cas9s used in genome editing (Methods). For these datasets, Protein2PAM achieved a median prediction accuracy of 0.797 (Fig. 2d). Utilizing the Protein2PAM confidence model removed 58 of 112 predictions (52%) but improved the overall median accuracy to 0.883 (Fig. 2d). Among the 79 diverse Cas9s characterized by Gasiunas et al. (3), Protein2PAM demonstrated the ability to rank its own predictions by their accuracy (Spearman’s r = 0.649, p = 1.03 × 10^-10^; Fig. 2e). Across proteins, Protein2PAM confidence scores exhibited a strong correlation with amino acid identity to training sequences (Spearman’s r = 0.804, p = 4.52 × 10^-19^), highlighting novelty as a key and interpretable factor in confidence estimation.

Overall, these results highlight the model’s robust performance for Type I and II systems and demonstrate its ability to match experimental outcomes despite being trained exclusively on evolutionary data.

In contrast, model performance was mixed for experimentally characterized Type V systems (Fig. 2d and Fig. S5). We tested Protein2PAM on 45 proteins from 11 Cas12 subtypes characterized in separate studies (Methods). Protein2PAM performed well for Cas12b and Cas12f (median accuracy = 0.772, *n* = 14) but was less accurate for Cas12a and other subtypes (median accuracy = 0.460, *n* = 31). For Cas12a in particular, the model tended to over-predict TTTN PAMs, which may be due to their high representation in the training dataset (Fig. S1). Overall, proteins from only three Cas12 subtypes were within 40% identity of any training sequence, highlighting the rarity of these systems in nature and underscoring the need for more training data.

Finally, we tested Protein2PAM on 20 engineered Cas9 and Cas12 proteins with altered PAM specificities from various studies (Methods). In most cases, the model predicted the same PAM as their wild-type counterpart with the exception of an Nme2Cas9 variant where Protein2PAM correctly predicted a shift from N_4_CC to N_4_CN (Fig. S6 and Table S2). Because Protein2PAM was trained on evolutionary data, the model may be insensitive to engineered mutations not observed in natural Cas proteins.

### Protein2PAM outperforms spacer-based PAM prediction

We compared the performance of Protein2PAM with PAMpredict (23), the most accurate bioinformatics tool for PAM prediction. Both tools were applied to predict PAMs for 11,381 Cas operons identified from genomic and metagenomic datasets not used for model training (Fig. 3a-b and Table S3). Protein2PAM predicted PAMs using protein sequences (Cas8, Cas9, and Cas12), while PAMpredict relied on CRISPR spacers aligned to a database of viral and plasmid genomes (21, 22). To enhance the sensitivity of Protein2PAM, we re-trained the models by integrating 157 experimentally characterized PAMs from the literature into the training dataset (Fig. 2d).

**Figure 3.**
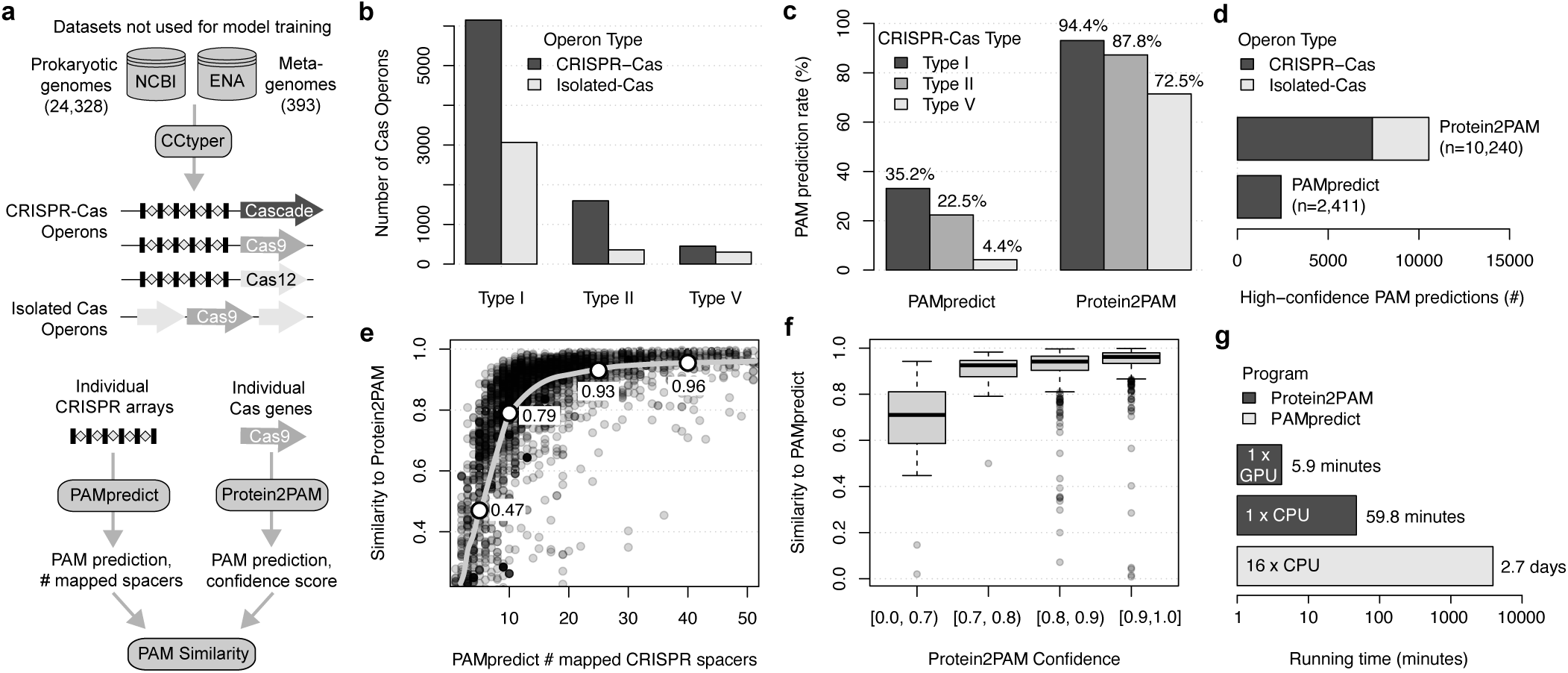
Rapid and sensitive PAM prediction with Protein2PAM. (a) We used CCtyper (32) to identify CRISPR-Cas operons in newly sequenced genomes and metagenomes. PAMs were predicted using PAMpredict from individual CRISPR arrays and Protein2PAM from individual Cas proteins. High-confidence predictions for PAMpredict required at least 10 mapped CRISPR spacers, while Protein2PAM high-confidence predictions were defined by confidence scores greater than 0.80. (b) Count of CRISPR-Cas operons and isolated Cas proteins (Cas8, Cas9, Cas12) identified by CCtyper. (c) Fraction of CRISPR-Cas operons with a high-confidence PAM prediction using PAMpredict and Protein2PAM. (d) Total number of high-confidence PAM predictions across methods for CRISPR-Cas operons and isolated Cas proteins, respectively. (e-f) PAM logos from Protein2PAM and PAMpredict were compared using the cosine similarity metric. PAMpredict predictions showed strong concordance with Protein2PAM when based on at least 10 mapped spacers, while Protein2PAM predictions with confidence scores >0.80 were highly concordant with PAMpredict. (g) Comparison of running times between Protein2PAM and PAMpredict.

Protein2PAM confidently predicted PAMs for 91.9% of 7,812 CRISPR-associated Cas operons, while PAMpredict yielded a confident prediction for only 30.9% (Fig. 3c). The largest difference was observed for Type V systems, where Protein2PAM was over 16 times more likely to yield a high-confidence prediction (72.5% vs. 4.4%) primarily due to insufficient spacer matches in the viral database using PAMpredict. Protein2PAM additionally provided predictions for Cas operons without associated CRISPR arrays, and overall produced 4.2 times more high-confidence predictions than PAMpredict (Fig. 3d). Protein2PAM predictions closely aligned with those of PAMpredict when both tools reported high-confidence in their respective predictions (Fig. 3e-f).

Lastly, we compared the running times and computational requirements of both approaches. PAMpredict was run on a Google Cloud instance with 16 vCPUs, taking 2.7 days to process 7,812 CRISPR-Cas operons (Fig. 3g). In contrast, Protein2PAM was run on a Google Cloud instance with one NVIDIA T4 GPU and completed the analysis of 11,381 Cas operons in just 5.9 minutes. Generating confidence scores extended Protein2PAM’s runtime to 59.8 minutes.

Together, these results demonstrate that Protein2PAM aligns with the current gold standard for PAM prediction, offers greater sensitivity, is independent of CRISPR spacer identification, and is considerably faster. For implementation details, refer to Data and Code Availability.

### *In silico* mutagenesis pinpoints protein-PAM interactions

To investigate whether Protein2PAM models have learned biophysical principles of PAM recognition, we performed *in silico* mutational scanning and identified point mutations predicted to alter PAM specificity (Fig. 4a, Table S4). Previous studies have identified PAM-interacting residues using Cas9 crystal structures bound to target DNA (29, 33–36) or experimental screening of Cas9 mutants (10, 11, 13, 37). In contrast, our models provide a computational alternative, enabling the identification of putative protein-DNA interactions across diverse Cas proteins without the need for structural or experimental data.

**Figure 4.**
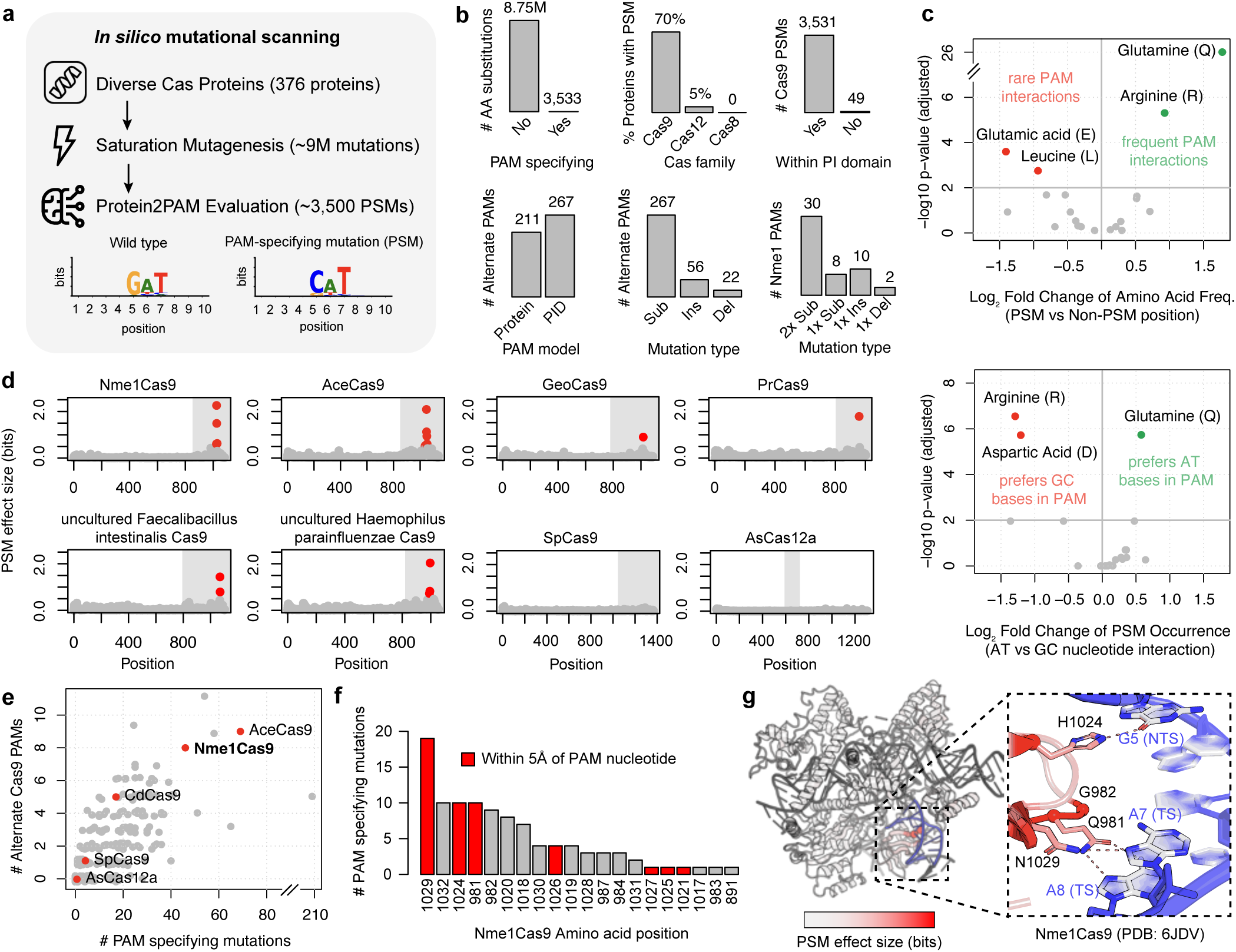
*In silico* mutagenesis pinpoints protein-PAM interactions. (a) Protein2PAM models were used to predict the effects of millions of single amino acid variants introduced into diverse Cas proteins. PAM-specifying mutations (PSMs) were defined as single residue changes that shifted the PAM by at least 0.5 bits at one or more nucleotide positions. (b) Barplots summarizing PSMs. While most mutations have no effect, most Cas9 proteins have a predicted PSM. Nearly all PSMs are located in the PI domain and are identified more sensitively using the PID-only Protein2PAM model. Substitutions are more effective than insertions and deletions at changing the PAM. Double amino acid variants expand the number of PSMs for Nme1Cas9. (c) Top: Volcano plot showing amino acids enriched at PSMs compared to their background distributions. Glutamine and arginine are notably overrepresented at PSMs. Bottom: Volcano plot showing the relative frequency of PSM interactions with AT versus GC nucleotides in the PAM. Glutamine PSMs preferentially interact with AT nucleotides, while arginine PSMs favor interactions with GC nucleotides. (d) Scatterplots depicting the distribution of mutational effects across eight Cas9 and Cas12 proteins. The y-axis indicates the maximum change in the PAM across all 20 mutations at each position. Shaded regions indicate PAM-interacting domains. (e) Scatterplot indicating the cumulative effect of single amino acid substitutions for different Cas proteins. Several proteins are predicted by Protein2PAM to be highly engineerable by single substitutions, including AceCas9 and Nme1Cas9. (f) The distribution of PSMs across amino acid positions for Nme1Cas9. PSM threshold reduced to 0.25 bits for barplot. (g) The protein structure of Nme1Cas9 superimposed with model predictions. Each amino acid position is colored by the maximum change in the PAM across all 20 mutations at the given position. Residues harboring PSMs are located in the PI domain and make hydrogen bonds with PAM DNA. Protein structure was visualized using PyMOL (38).

To comprehensively map the landscape of PAM interactions in Cas9, we used the full-sequence Type II PAM model to predict the effects of over 8 million single amino acid substitutions across 336 phylogenetically diverse Cas9s, including 15 previously applied in genome editing. We defined PAM-specifying mutations (PSMs) as those predicted to alter PAM specificity, measured by an L1 distance shift of ≥ 0.5 bits at one or more PAM nucleotide positions (Fig. 4a).

The vast majority of mutations were predicted to have no effect on PAM recognition, with only 0.04% classified as PSMs (Fig. 4b). Strikingly, large-effect mutations clustered within the PI domain and stood out as clear outliers relative to neighboring sites (Fig. 4d). Notably, 99.98% of PSMs were located within the annotated PI domain, even though this region accounted for only 23.7% of the cumulative protein length. These results suggest that the full-sequence Cas9 model relies almost exclusively on the PI domain, reaffirming its critical role in determining PAM specificity.

In contrast to Cas9, point mutations in Cas8 and Cas12 had minimal impact on the predicted PAM (Fig. 4b). After testing over 500,000 mutations across 40 phylogenetically diverse Cas8 and Cas12 proteins, we predicted only two PSMs. However, both mutations were located at the same amino acid position and were just above the threshold for classifying a mutation as a PSM. By contrast, we found PSMs for 70% of the 336 Cas9 proteins we analyzed. These findings highlight that PAM customization with Protein2PAM is limited for Type I and V systems due to PAM conservation (Fig. S1) but likely effective for Type II systems.

To obtain higher resolution of protein-PAM interactions, we applied the PAM model trained specifically on Cas9’s PAM-interacting domain (PID-only model). Compared to the full-sequence model, the PID-only model predicted 12.9% more PSMs and 30.5% more alternate PAMs (Fig. 4b). Next, we used this model to test all possible single amino acid insertions and deletions within the PI domain. However, indels were less effective than substitutions for PAM diversification (Fig. 4b) and were predicted to result in reduced enzyme fitness (Fig. S7). Interestingly, we identified several Cas9 orthologs that appeared amenable to engineering with single substitutions – point mutations resulted in at least eight alternate PAMs for previously characterized Cas9s like Nme1Cas9 (39) and AceCas9 (40), as well as for five novel Cas9s identified from human microbiome samples (Fig. 4e). We expect further computational screening to uncover additional Cas9 orthologs with broad potential for PAM engineering with Protein2PAM.

Next, we examined whether any patterns emerged among the predicted PSMs. Several amino acids were highly overrepresented among PSMs (Fig. 4c), such as glutamine and arginine, which were 3.4x and 1.9x more likely to be found at a PSM position compared to the rest of the PI domain (*X*^2^, *q*-values *<* 5 × 10^−6^). We also observed strong preferences between these amino acids and specific nucleotides (Fig. 4c), consistent with previously observed amino-acid nucleotide interactions (41), including the propensity for glutamine in Cas9 to recognize adenine in the major groove of DNA (29).

Finally, we analyzed the locations of PSMs in crystal structures of eight Cas9 proteins. (33, 34). Strikingly, many top-ranked mutations identified by Protein2PAM occurred at residues forming sequence-specific contacts with PAM DNA (Fig. S8). In total, 58.5% of the 159 identified PSMs in these proteins were located at residues that form hydrogen bonds with PAM nucleotides (*X*^2^, *p*-value *<* 2.2×10^−16^), and this percentage increased to 80.0% when considering only the 50 PSMs with the largest effect. Notably, PSMs were not found at all PAM-interacting positions. For example, in SpCas9, Arg1333 forms a critical interaction with the guanine at the second PAM position (NGG), but due to its high conservation in nature, mutations were not predicted to alter PAM recognition. Overall, these findings suggest that our evolutionary-informed models have captured key biophysical interactions governing protein-to-PAM recognition across diverse Cas9 proteins.

### Computational evolution of PAM-customized Nme1Cas9 variants

We hypothesized that Protein2PAM models could be used to generate PAM-customized enzyme variants. To test this hypothesis, we focused on Nme1Cas9, for which single amino acid mutations yielded several alternate PAMs (Fig. 4e) and high PAM diversity has been observed among closely related orthologs in nature (31). We initially selected four Nme1Cas9 point mutations (N1029A, Q981A, H1024D, and H1024E) predicted to induce large shifts in PAM specificity and produce distinct PAMs (Fig. 4f). Enzyme variants were experimentally characterized using the high-throughput PAM determination assay (HT-PAMDA) in human cell lysate (8, 42), which measured Cas9-mediated depletion of target sequences flanked by a library of all possible PAMs (Fig. 5a). Deep sequencing at four time points was used to calculate the cleavage rates for each enzyme on a library of substrates encoding all possible PAMs (Table S8).

**Figure 5.**
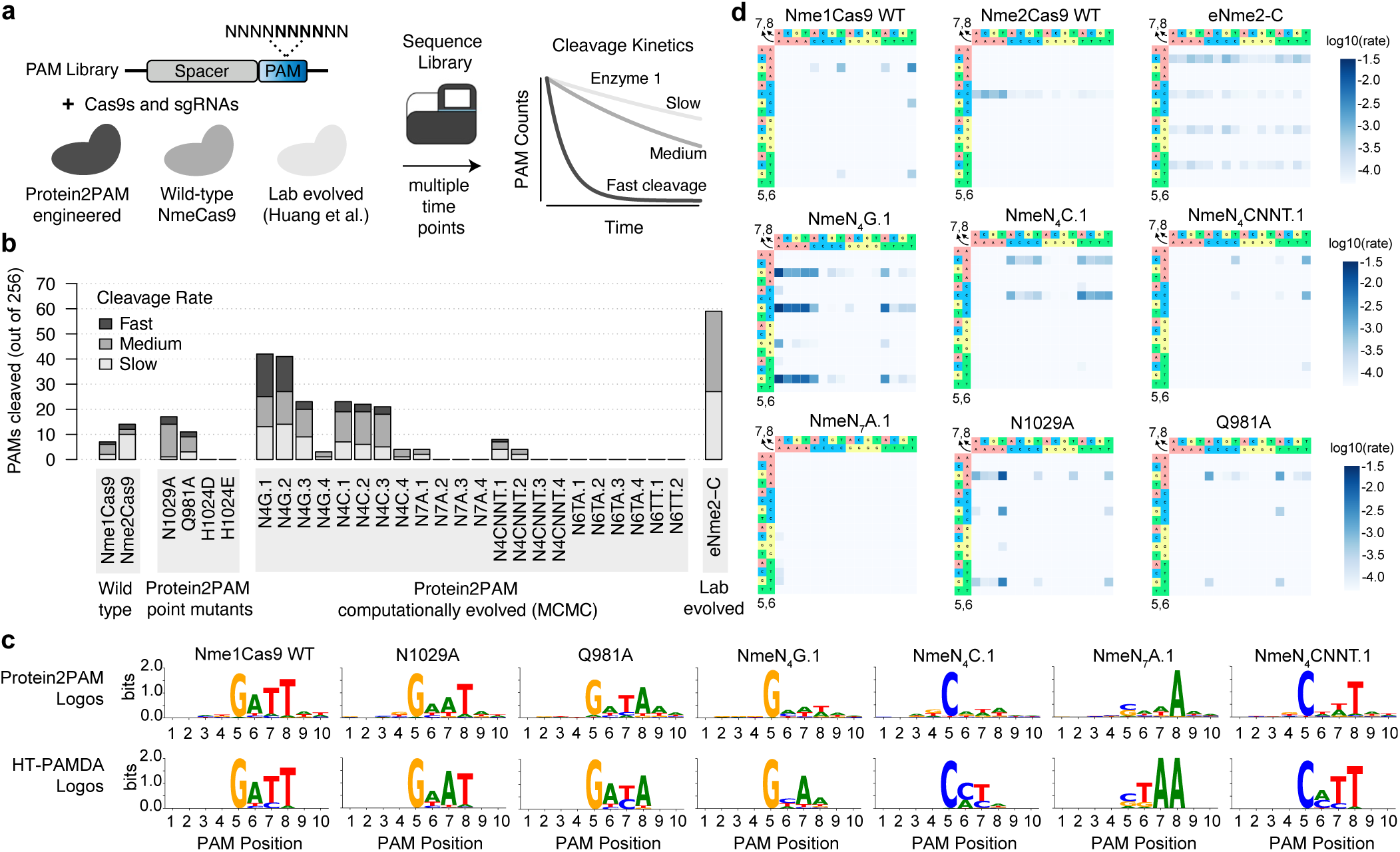
Computational evolution of PAM-customized NmeCas9 variants. (a) Proteins were characterized using the high-throughput PAM determination assay (HT-PAMDA) in human cell lysate, measuring Cas9 cleavage rates on substrates with all possible PAMs. Cleavage rates were quantified at positions 5 to 8 of the PAM library after deep sequencing at four time points. (b) Activity landscape across Nme1Cas9 enzyme variants. The cleavage rate of each PAM is derived by tracking depletion over four time points (Fast: rate > 1e-3, Medium: rate > 1e-4, Slow: rate > 5e-5). (c) Top: PAM logos predicted using Protein2PAM. Bottom: PAM logos generated from HT-PAMDA data. For HT-PAMDA logos, each four-nucleotide PAM was weighted by its corresponding rate constant, nucleotide counts were normalized to frequencies summing to 1.0 per position, and frequencies were converted to information content. (d) HT-PAMDA heatmaps which display rate constants for different enzyme variants at PAM positions 5-8.

Two of the four mutants (N1029A and Q981A) exhibited robust cleavage along with a shift in PAM preferences that closely aligned with model predictions (Fig. 5b-d). N1029A and Q981A cleaved their predicted PAMs — N_4_GNAT and N_4_GNTA — at rates 270x and 9.4x higher than wild-type Nme1Cas9 (Fig. 5d and Table S5), while showing 4.7x and 13.9x lower cleavage rates at the preferred PAM of Nme1Cas9 (N_4_GATT). Examining Nme1Cas9’s crystal structure, the side chains of N1029 and Q981 form base contacts with PAM nucleotides 7 and 8 (Fig. 4g), while Protein2PAM predicted that mutations at these residues would result in shifts at the corresponding PAM nucleotides. In contrast, the two H1024 mutations abolishe activity in the HT-PAMDA assay, even though H1024D is observed in Nme orthologs that recognize N_4_C PAMs (31). While H1024 is involved in PAM recognition (33), this single mutation alone appears insufficient for recognizing N_4_C PAMs.

Next, we harnessed Protein2PAM models to design variants for PAMs not achievable by single substitutions. While Nme1Cas9 has been used for genome editing due to its small size and high specificity (39), its long PAM (N_4_GATT) limits the number of editable sites in the human genome. Our objective was to use Protein2PAM to engineer Nme1Cas9 variants with broader PAM compatibility, specifically targeting three single-nucleotide PAMs (N_4_G, N_4_C, N_7_A) and three di-nucleotide PAMs (N_6_TT, N_6_TA, N_4_CNNT). These PAMs were chosen after examining nucleotide conservation patterns across PAMs from Nme orthologs.

To computationally evolve Nme1, we employed the Gibbs with Gradients Markov Chain Monte Carlo (MCMC) algorithm (43), iteratively introducing mutations within the PI domain to guide each protein variant toward its target PAM while maintaining fitness, as estimated by ProGen2 (15). To increase sensitivity, we trained and utilized a variant of the Protein2PAM model in which Nme1Cas9 orthologs were upweighted, and to preserve enzymatic function, we sampled candidate mutations from a multiple sequence alignment of closely related Nme orthologs. MCMC trajectories were terminated after 2000 steps, at which point most had converged, and we considered variants at all points along the trajectories.

We used our pipeline to run 30 trajectories per PAM, generating 30,000 Nme1Cas9 variant enzymes, and selected 22 variants targeting the six PAMs for experimental characterization (N_4_G, N_4_C, N_7_A, N_6_TT, N_6_TA, N_4_CNNT). Variants contained an average of 11.6 mutations (range: 5–18) and were selected based on their predicted similarity to the target PAM, pLM log-likelihood, and mutation count. Engineered variants were experimentally characterized using HT-PAMDA with two guide RNAs (gRNA), alongside Nme1Cas9 and Nme2Cas9 wild-type enzymes (Fig. S9). An enzyme was deemed active in the HT-PAMDA assay if it cleaved at least one PAM with a rate constant (*k*) > 5 × 10^−5^ (Methods).

Strikingly, among the 22 tested enzymes, 50% exhibited activity, with 6 showing cleavage rates surpassing those of the wild-type enzymes. (Fig. 5b). Across design targets, a high fraction of the sequences were active for N_4_G, N_4_C, and N_4_CNNT PAMs (10/12), while few enzymes were active for other target PAMs, including N_7_A, N_6_TT, and N_6_TA (1/10). This indicates that despite an overall high hit rate, certain PAMs were difficult to achieve using Protein2PAM. Sequence logos generated from HT-PAMDA data for active enzymes generally aligned with model predictions, though there was some evidence of overfitting to sequences generated during MCMC (Fig. S10).

The most active variant was designed for N_4_G PAMs and contained 13 mutations relative to Nme1Cas9 (D957G, V979I, V980K, Q981A, Q989T, N996E, S1000V, M1016K, G1018A, N1029A, N1031S, I1041V, E1048Q). We named this enzyme NmeN_4_G.1. NmeN_4_G.1 had a considerably broadened PAM, exhibiting cleavage at 42 N_4_G PAMs, compared to just 7 for Nme1Cas9 (Fig. 5b). It also demonstrated significantly higher peak activity, cleaving its top 10 PAMs 56.4x faster than Nme1Cas9’s top 10 PAMs. However, NmeN_4_G.1 did display a preference for A at position 7 of the PAM (N_4_GNAN) suggesting that further optimization is required to fully meet the design goal (Fig. 5c-d). Our most active enzyme designed for N_4_C PAMs, NmeN_4_C.1, also showed a clear shift in specificity towards its design target, cleaving 21 N_4_C PAMs compared to 14 for Nme2Cas9 and 0 for Nme1Cas9. NmeN_4_C.1 also exhibited enhanced peak activity, with its top 10 PAMs cleaved 9.6x faster than Nme2Cas9’s top 10. Notably, all N_4_C designed enzymes contained the H1024D mutation, despite the H1024D single point mutant being completely inactive.

Next, we compared the NmeN_4_G.1 and NmeN_4_C.1 variants to eNme2-C, a broad-PAM Nme2Cas9 variant engineered over multiple rounds of phage-assisted directed evolution to yield a N_4_C PAM (13). While eNme2-C cleaved a greater number of PAMs (*n* = 59) compared to NmeN_4_G.1 (*n* = 42) and NmeN_4_C.1(*n* = 21), it exhibited significantly slower kinetics in the HT-PAMDA assay. eNme2-C cleaved its top 10 PAMs 21.5x slower than NmeN_4_G.1 and 2.7x slower than NmeN_4_C.1. We also tested the eNme2-T.1 PAM-engineered variant (13), but the enzyme displayed no activity in HT-PAMDA. These results highlight the ability of Protein2PAM to efficiently engineer Cas enzymes with novel PAMs and enhanced activity without the need for experimental training data, iterative screening, or structural modeling.

Having achieved customization for specific, user-defined PAMs, we aimed to design enzyme variants that maximized PAM diversity. To this end, we adopted a new design strategy, generating 37 million variants with up to 5 combinations of 102 PAM-specifying mutations (PSMs). To broaden PAM diversity, we relaxed constraints by allowing mutations beyond those observed in Nme1 orthologs and lowered the threshold for defining a PSM to 0.25 bits. For experimental validation, we selected 178 variant enzymes, each containing 1 to 5 PSMs which targeted 64 alternate PAMs. Enzymes were selected to maximize pLM log likelihoods and minimize mutation count.

In contrast to our computationally evolved enzymes, the combinatorial mutants showed a markedly lower success rate, despite having fewer mutations and higher pLM log likelihoods (Fig. S11). Among the 178 tested enzymes, only 18 were active (10.1%), and just 8 displayed cleavage rates exceeding those of the wild-type enzymes (4.5%). Notably, a significant fraction of enzymes contained one or more “non-natural” mutations absent from closely related Nme1 orthologs. Activity was substantially lower for these enzymes (12 out of 166) compared to those containing only natural mutations (6 out of 11). These findings suggest that sampling from natural mutation distributions may be crucial for achieving effective single-shot PAM customization with Protein2PAM.

## Discussion

Protein2PAM is a protein language model that efficiently predicts the PAM specificity of CRISPR-Cas systems directly from Cas protein sequences. We demonstrated that Protein2PAM accurately predicts the PAMs of naturally occurring proteins and identifies PAM-interacting residues without relying on structural information. Using Protein2PAM, we computationally evolved Nme1Cas9 enzyme variants that, through experimental validation with HT-PAMDA, exhibited both higher activity and broadened PAM specificity compared to wild-type enzymes. This work represents the first successful demonstration of machine learning being used to precisely alter DNA recognition in Cas enzymes in a single step and without relying on laboratory training data.

While powerful, current Protein2PAM models have several limitations. Most notably, they are constrained by the natural co-variation of Cas proteins and their PAMs. PAMs for Type II systems were found to rapidly shift over evolution, enabling Protein2PAM to learn which residues in Cas9 are important for PAM specificity. However, the high conservation of PAMs in Type I and Type V systems limits protein-PAM covariation, making the current models unsuitable for engineering these systems. Using HT-PAMDA, we experimentally validated the model’s capability to guide the engineering of PAM-customized Nme1Cas9 variants. However, this process may be more challenging for other Cas9 orthologs with fewer training examples or reduced protein-PAM co-variation. Finally, Protein2PAM models do not account for protein fitness, sometimes predicting alternate PAMs for enzymatically inactive mutants. To address this, it will be important to couple Protein2PAM with methods that reliably predict mutational fitness within the hypervariable PAM-interacting domain.

We see several promising future directions. With the exponential growth of genomic databases, we aim to automate model updates, enabling Protein2PAM to evolve alongside data growth and continually enhance its understanding of protein-PAM interactions across the tree of life. Furthermore, our models have the potential to incorporate data from experimental screening of enzyme variants, which could establish a feedback loop to optimize Protein2PAM for more efficient protein engineering. We also envision exciting applications of Protein2PAM, including engineering a library of PAM-specific enzyme variants capable of targeting any site in the human genome and optimizing Cas9 variants for specific therapeutic targets. Finally, our framework could be adapted for other DNA-binding proteins, such as recombinases, transcription factors, and zinc fingers, paving the way for machine learning to precisely tailor diverse DNA-binding proteins for therapeutic applications.

## Methods

### Curation of CRISPR-Cas sequences

Cas proteins and their associated CRISPR arrays were identified from the CRISPR-Cas Atlas, as previously described (17). The resource contains 1,246,163 CRISPR-Cas operons that were derived from 26.2 Tbp of genome and metagenomic assemblies. For modeling PAMs, we focused on a subset of 653,991 operons from CRISPR Types I, II, and V where we could confidently identify an effector protein linked to a CRISPR array.

For Cas9 proteins, we also identified PI domains using a custom-built database of 123 profile HMMs. PI domain sequences were sourced for 9,161 diverse proteins (3), de-replicated at 90% identity using CD-HIT v4.8.1 (44), aligned using DIAMOND v2.1.6 (options: –query-cover 80 –subject-cover 80 –very-sensitive) (45), and clustered using MCL v22.282 (options: -I 1.5) (46). Multiple sequence alignments (MSAs) were created with FAMSA v2.2.2 (47) and used as input to hmmbuild v3.4 (48). HMMs were aligned to Cas9s from the CRISPR-Cas Atlas using hmmsearch with a 1e-5 E-value threshold. For proteins lacking a valid PI domain alignment, we instead extracted the region downstream of RuvC III based on alignment to the RuvC Pfam domain (PF18541).

### Bioinformatic PAM determination

PAMs for CRISPR-Cas systems were characterized by aligning CRISPR spacers to viral and plasmid genomes and performing statistical analysis of regions flanking protospacers. To enhance the number of spacers associated with each Cas ortholog, we pooled CRISPR arrays from closely related Cas proteins. Cas proteins included Cas8 for Type I systems, Cas9 for Type II systems, and Cas12 for Type V systems. Proteins were clustered using MMseqs2 13.45111 with default parameters (25) at 100%, 99%, and 98% amino acid identity (see below for details).

Each pool of spacers contained CRISPR arrays in varying orientations. To address this, CRISPR repeats associated with each Cas protein cluster were aligned using CD-HIT (options: cd-hit-est -c 0.95 -s 1.0) (44) and CRISPR spacers were consistently oriented based on the orientation of aligned repeats. CD-HIT was also used to de-replicate CRISPR spacers within each cluster to minimize the impact of overrepresented sequences (cd-hit-est -c 0.90 -T 1 -s 0.90).

Oriented and de-replicated pools of CRISPR spacers were input to PAMpredict v1.0.2 (23). This tool aligned spacers to a database of 16 million virus and plasmids genomes from IMG/VR v4 (21) and IMG/PR (22), extracted 10-nt protospacer flanking regions, computed nucleotide frequencies, and identified sequence motifs. PAMs were detected upstream of protospacers for Type I and V systems and downstream for Type II systems. The strand of the PAM was determined based on the 10-nt region containing a more conserved DNA motif. A PAM was classified as high-confidence based on two criteria. First, it needed to be identified from at least 10 unique protospacers, following the recommendation of Ciciani et al. (23). Second, we required a signal-to-noise ratio greater than 2.0 (Fig. S1). For Type II systems, the signal-to-noise ratio was calculated as the ratio of the maximum information content across the 10 nucleotide positions upstream and downstream of the protospacer, and conversely, for Type I and Type V systems, the ratio was calculated in the opposite direction.

Each Cas protein was associated with multiple PAM predictions due to the varying MMseqs2 clustering thresholds. Clustering at lower identity thresholds increases the number of CRISPR spacers linked to a protein, improving the likelihood of PAM detection, but also increasing the chances of pooling Cas variants with different PAM specificities. To mitigate this, we selected the PAM prediction at the highest percent identity clustering threshold that met our prediction quality criteria. We compared our PAM dataset to two previously published studies. In the study by Ciciani et al., PAMs were bioinformatically quantified for Cas9 proteins clustered at 98% amino acid identity. Using this threshold, Ciciani et al. identified PAMs for 2,546 Cas9 protein clusters with at least 10 mapped spacers, whereas our study reported PAMs for 7,229 Cas9s clustered at 98% identity (2.8x increase). Similarly, Gasiunas et al. experimentally characterized PAMs for 79 unique Cas9 proteins, compared to the 15,731 unique Cas9 proteins with bioinformatically characterized PAMs in our study (199x increase).

### Training the Protein2PAM models

Both the PAM prediction and PAM confidence models consist of a 650 million parameter transformer encoder (16) with an MLP head, which has one hidden layer with embedding dimension 1280 (matching that of the transformer encoder). In all cases, we evaluated our models using 10-fold cross-validation and ensured that the validation data came from different 90% identity clusters from the training data.

For Protein2PAM, the MLP head takes as input the [*CLS*] embedding vector from the transformer encoder and has an output dimension of 40. The output is reshaped into a 10x4 matrix and transformed into a sequence of probability distributions over nucleotides with a softmax (Fig. 2a). The transformer encoder was initialized with the pretrained ESM-2 model, but its weights received gradient updates during training. We trained each model to maximize the sum of the negative cross entropy and PAM similarity between true and predicted PAMs, using PyTorch Distributed Data Parallel on machines with 2 A100 GPUs. Each training batch contained up to 2500 tokens, and we accumulated the gradient for 4 steps before updating model weights. We used the Adam optimizer with a learning rate of 0.0001 (all other hyperparameters set to PyTorch defaults). Training was stopped when the validation loss did not improve for 5000 steps, and we used the checkpoints with the best validation loss. See Table S9 for a full list of different Protein2PAM models and manuscript analyses they are associated with.

### Estimating Protein2PAM prediction confidence

For Protein2PAM confidence estimation, we first calculated the percent identity between the input sequence and its 10 nearest neighbors in the training data. These 10 percent identities were encoded into a 200-dimensional vector using piecewise linear embeddings (PLE), which was then projected into a 1280-dimensional space via a linear layer. This vector was added to the 1280-dimensional [*CLS*] embedding from the transformer encoder before passing the combined vector through a 2-layer MLP. A sigmoid activation was applied to the MLP output, constraining it to the [0, 1] range, where it could be interpreted as the predicted PAM similarity (Fig. 2b).

For each CRISPR-Cas type, we first trained a CasEncoder, a 650-million parameter transformer initialized from a pretrained ESM-2 checkpoint. The CasEncoder was fine-tuned using the masked language modeling loss on proteins from the CRISPR-Cas database to learn a consistent representation of the relevant protein family. Once trained, the CasEncoder weights were frozen, and the proteins were encoded using their [*CLS*] token embeddings. We computed the percent identity between each sequence and the 10 most similar sequences in the training dataset, embedding these values using PLE and combining them with the CasEncoder embeddings.

The combined embeddings were passed through a 2-layer MLP, which was trained by minimizing the mean squared error between the predicted PAM similarity and the accuracy of Protein2PAM’s prediction. We used the Adam optimizer with a learning rate of 0.0003 and a batch size of 1024. The best performing confidence model was selected based on the checkpoint with the lowest validation loss.

### Quantifying PAM similarity

We quantified the similarity between two PAMs based on their information content rather than probability distributions. Information content is measured using the relative entropy between *P* and a background distribution *Q*, where *Q* is uniformly distributed across *A*, *C*, *G*, and *T*. Specifically, the information content of nucleotide *n* at position *i* is calculated as: 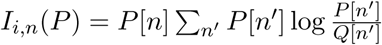.

Given two 10 × 4 PAM information matrices, *I*^(1)^ and *I*^(2)^, the cosine similarity between their vectorized forms provides a natural similarity metric. However, this fails to distinguish between positions where one PAM has low information (denoted as *N*) and the other has high information. To address this, we augmented each position in the matrix with the information content of a fictitious *N* nucleotide. This *N* content is high when the original PAM has low information at that position, but the comparison PAM has high information, and low when both PAMs have either high or low information.

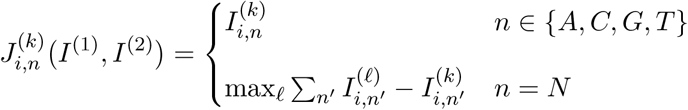

Finally, we computed the cosine similarity between the vectorized forms of the augmented information matrices, *J* ^(1)^ and *J* ^(2)^, to obtain the PAM similarity. This augmented similarity metric is used to determine accuracy when comparing to a ground truth PAM and is referenced throughout this paper.

### Benchmarking on experimental datasets

Protein2PAM models were evaluated on experimentally determined PAMs for diverse CRISPR systems (Table S1). For Type I systems, Protein2PAM was applied to 14 Cas8 proteins with characterized PAMs (30). For Type II systems, Protein2PAM was applied to 79 Cas9 proteins spanning the phylogeny (3), 23 Cas9 proteins from closely related Type II-C systems (31) and 10 Cas9 proteins used as genome editors, including: SpCas9, St1Cas9 and St3Cas9 (2), Nme1Cas9 and Nme2Cas9 (49), AceCas9 (50), FnCas9 (51), FrCas9 (52), CjCas9 (53), and CdCas9 (54). For Type V systems, Protein2PAM was applied to 45 Cas12s with experimentally characterized PAMs, including: Cas12a (55, 56), Cas12b (57), Cas12d (58, 59), Cas12f (60), Cas12h and Cas12i (4), Cas12j (61), Cas12k (62), Cas12l (63), Cas12m (64), and Cas-lambda (65). Lastly, Protein2PAM was applied to 20 engineered proteins from the literature with altered PAM specificities, including variants of: SpCas9 (7, 8, 11), SaCas9 (10), St1Cas9 (66), Nme2Cas9 (13), CjCas9 (37), and Cas12a (9, 67, 68).

### *In silico* mutational scanning

We performed a large-scale mutagenesis experiment to identify point mutations predicted to change the PAM. Diverse wild-type proteins were selected for Cas8, Cas9, and Cas12 from 70% identity protein clusters from the CRISPR-Cas Atlas training dataset. For Cas9, we selected proteins from 336 clusters containing at least 20 members. These were supplemented with 15 Cas9s from the literature that have been used in genome editing and include: AceCas9, CdCas9, CjCas9, FnCas9, FrCas9, GeoCas9, Nme1Cas9, Nme2Cas9, PrCas9, SaCas9, ScCas9, SpCas9, St1Cas9, St3Cas9, and TnCas9. For Cas8 and Cas12, we selected proteins from the top-20 largest clusters. We generated all possible single amino acid variants – including substitutions, insertions, and deletions – from each wild-type protein sequence. For Nme1Cas9, we additionally generated all possible double amino acid substitution variants. Cas8, Cas9, and Cas12 wild-type and mutant proteins were used as input to the corresponding full-sequence Protein2PAM models. For Cas9, annotated PI domain regions were also used as input to Protein2PAM PID-only Cas9 model.

To evaluate the impact of mutations on PAM specificity, we compared the Protein2PAM-predicted PAM profiles for wild-type sequences and their corresponding single amino acid mutants. The effect size of each mutation was quantified using the maximum *L*_1_ distance across the 10 PAM positions, defined as: 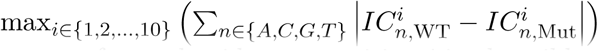, where 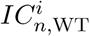 and 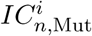 represent the information content for nucleotide *n* at position *i* in the wild-type and mutant sequences, respectively. PAM-specifying mutations (PSMs) were classified if they caused a measurable shift in PAM specificity, indicated by an *L*_1_ distance change of ≥ 0.5 bits at one or more nucleotide positions in the predicted PAM.

### Design of PAM-customized Nme1Cas9 variants

To design proteins that targeted specific PAMs, we leveraged the Gibbs with Gradients Markov Chain Monte Carlo (MCMC) algorithm (43). MCMC provides a stochastic method that iteratively introduces *in silico* mutations to a protein sequence that are expected to improve its score according to an oracle model. We averaged two components to compute a score for a protein sequence: the Protein2PAM loss between the predicted PAM and a target PAM, and the language modeling loss of ProGen2 (15) fine-tuned on the CRISPR-Cas Atlas (17). To increase sensitivity, we trained and utilized a variant of the Protein2PAM model where NmeCas9 orthologs were upweighted in the training data. To preserve enzymatic function, we only sampled candidate mutations in the PI domain (positions: 937–1082) from a multiple sequence alignment of NmeCas9 orthologs that had at least 70% identity to Nme1Cas9. We ran all MCMC trajectories for 2000 steps. For each target PAM, we selected the variants at any point along the trajectories which individually minimized the Protein2PAM loss, the fine-tuned ProGen2 model’s loss, and the aggregate score.

To design proteins targeting diverse PAMs, we adopted a combinatorial mutagenesis approach. We first identified a minimal set of 102 PSMs that shifted Nme1Cas9’s PAM preference by at least 0.25 bits. Notably, 73 of these mutations were concentrated at seven key sites, including three (Q981, H1024, N1029) that form hydrogen bonds with PAM DNA in Nme1Cas9’s WT structure. Pairwise combinations of these 102 mutations yielded 9,464 double, 132,838 triple, 2,530,861 quadruple, and 34,777,000 quintuple mutants which were predicted to collectively target 177 alternative PAMs. For experimental validation, we selected 178 of these variants, each containing 1 to 5 PSMs which targeted 64 alternate PAMs. Enzymes were selected to maximize PAM diversity, minimize mutation count, and maximize pLM log likelihoods as measured by ProGen2 fine-tuned on the CRISPR-Cas Atlas. All NmeCas9 variants are listed in Table S5.

### Plasmid construction and gRNA *in vitro* transcription

pCMV-Nme1Cas9-P2A-EGFP was synthesized by Twist Biosciences and designed to harbor the wild-type Nme1Cas9 sequence and serve as our expression plasmid (Table S6). The PAM-interacting domain regions of computationally designed enzymes were codon-optimized for *Homo sapiens* and synthesized by Twist Biosciences. DNA was ordered as an arrayed, lyophilized, double-stranded DNA fragment library. DNA fragments contained two flanking regions with complementary overlap to the wild-type WED domain (57 bp complementarity) and the pCMV plasmid (30 bp complementarity) for downstream cloning. To generate the arrayed plasmid variant library, the expression plasmid was first linearized with inverse PCR to remove the wild-type PID with the following recipe: 25 µL 2x Platinum SuperFi II PCR master mix (Invitrogen), 1.25 µL 10 µM forward primer, 1.25 µL 10 µM reverse primer, 10 ng of template, and nuclease-free water to a final volume of 25 µL per reaction. The following PCR parameters were then applied: initial denaturation at 98C for 30 s, followed by 15 cycles of denaturation at 98C for 10 s, annealing at 60C for 30 s, extension at 72C for 4.2 min, and a final extension at 72C for 5 min. The PCR linearized backbone was then incubated with DpnI (NEB) at 37C for 1 hr to digest the residual template.

PID variant fragments were resuspended to a concentration of 20 ng/µL in 50 µL of 10 mM Tris-Cl, pH 8.5, then introduced into the linearized expression plasmid through HiFi assembly (NEB) with a 10:1 insert-to-vector ratio. Reactions were incubated at 50C for 20 min, then transformed into NEB^®^ Turbo Competent *E. coli* cells. Colony PCR was performed to screen clones for proper assembly (REDTaq^®^ DNA Polymerase Master Mix, VWR), and passing clones were mini-prepped (Qiagen) and validated with whole plasmid sequencing (Table S6).

The two plasmid libraries encoding 10-nt randomized PAMs and different spacer sequences were generated similar to previously described (8, 42). Briefly, the plasmid p11-lacY-wtx1 (69) (Addgene ID 69056) was digested with EcoRI-HF, SpeI-HF, and SphI-HF (NEB) and purified. The PAM libraries were generated by annealing oNK507 or oNK508 with oBK984 (Table S7) and performing an extension reaction with Klenow fragment (3’ to 5’ exo-) (NEB) prior to digestion with EcoRI-HF. The digested duplexed libraries were ligated into the digested p11-lacY-wtx1 backbone, with the ligations cleaned up and transformed into XL1-Blue electrocompetent cells. The resulting transformation was grown overnight and maxiprepped.

For *in vitro* transcription of gRNAs, the pT7-SpCas9-sgRNA-scaffold plasmid (MSP3485; Addgene ID 140082) was digested with NheI-HF and HindIII-HF (NEB) to remove the T7 promoter and SpCas9 gRNA scaffold. Pairs of oligonucleotides encoding the T7 promoter, spacer sequences, and Nme1Cas9 gRNA scaffold (Table S7) were annealed and ligated into the digested MSP3485 plasmid to generate the final gRNA IVT plasmids (Table S6). gRNA transcription reactions were performed by digesting these IVT template plasmids with DraI, utilizing 10 µL of digested plasmid as template in reactions from the T7 RiboMAX Express Large Scale RNA Production System kit (Promega) for 19 hours at 37C, cleaning up the IVT reactions via MinElute PCR Purification Kit (QIAGEN), and quantifying gRNA yield.

### HT-PAMDA screening

HEK 293T cells (ATCC) were maintained at 37 °C and 5% CO2 in DMEM (Gibco) supplemented with 10% FBS (Gibco) and 1% penicillin/streptomycin. For mycoplasma testing, supernatant media was analyzed via PCR.

To generate human cell lysates containing Nme1Cas9 variant enzymes, HEK 293T cells were seeded at a density of ∼1.5 × 10^5^ cells per well in a 24-well plate ∼20 hours prior to transfection, and transfected with a mixture of approximately 800 ng of nuclease-P2A-EGFP expression plasmid (Table S6) and 1.5 *µ*l of TransIT-X2 (Mirus) in a total of 50 *µ*l of Opti-MEM (ThermoFisher). After 48 hours following transfection, cells were lysed using 100 *µ*l of lysis buffer [final concentration of 20 mM Hepes, pH 7.5; 100 mM KCl; 5 mM MgCl_2_; 5% (vol/vol) glycerol; 1 mM DTT; 0.1% (vol/vol) Triton X-100; and 1x SigmaFast Protease Inhibitor Cocktail tablet (EDTA-free)] and normalized using a DTX 880 Multimode Plate Reader (Beckman Coulter) based on EGFP fluorescence to 150 nM Fluorescein (Sigma).

HT-PAMDA reactions were performed similar to previously described (8, 42). Briefly, the PAM library plasmids were linearized with PvuI-HF (NEB). For each HT-PAMDA reaction, 5.625 *µ*l of normalized cell lysate and 4.5 *µ*l of *in vitro* transcribed 2.5 *µ*M gRNAs were incubated at 37 °C for 10 min to pre-form the Cas9 RNPs, and the buffer and 56.25 fmols of linearized PAM library plasmid were added to initiate *in vitro* cleavage reactions. The cleavage reactions were performed at 37 °C, and 5 *µ*l were removed at 1 min, 8 min, 32 min, and 135 min and terminated by adding 5 *µ*l of stop buffer (50 mM EDTA and 2 mg/ml proteinase K (NEB)).

The remaining uncleaved PAM library from each reaction time point for each Nme1Cas9 variant were PCR amplified with Q5 polymerase (NEB) using cycling conditions of 98 °C for 2 min, 30 cycles of 98 °C for 10 s, 67 °C for 10 s and 72 °C for 10 s, and 72 °C for 1 min, and the PCR primers encoding 5 nt barcodes (Table S7). For each time point, PCR products were pooled, purified with paramagnetic beads (prepared as previously described (9, 70) and PCR amplified using primers encoding Illumina i5 and i7 indexes (Table S7) using cycling conditions of 98 °C for 2 min, 10 cycles of 98 °C 10 s, 65 °C for 30 s and 72 °C for 30 s, and 72 °C for 5 min. The libraries were sequenced on a NovaSeq X Plus (Illumina).

### Analysis of HT-PAMDA data

The sequencing results were analyzed using a modified version of the HT-PAMDA analysis scripts, available at: https://github.com/RachelSilverstein/HT-PAMDA-2. HT-PAMDA results for all enzymes analyzed via this method are available in Table S8. For each enzyme tested, the pipeline calculated cleavage rate constants (*k*) for all four-nucleotide PAMs in the library (positions 5–8), using two distinct spacer sequences. Rate constants for each PAM were averaged across spacer sequences. We observed low technical variability between spacer sequences, with a mean *r*^2^ = 0.976 when comparing PAM cleavage rates between the gRNAs for each active enzyme.

We categorized cleavage rates as follows: High (*k >* 10^−3^), Medium (10^−3^ ≥ *k >* 10^−4^), and Slow (5 × 10^−5^ ≤ *k <* 10^−4^) (Fig. 5a). These thresholds were chosen based on the PAM cleavage rate distribution for wild-type Nme1Cas9, for which only one PAM (N_4_GATT) exceeded the High cleavage rate threshold (*k* = 3.4 × 10^−3^) and cleavage rates were categorized as Slow for 97% of PAMs. To summarize activity, we counted the number of PAMs at each activity level for each enzyme (Fig. 5b).

To compare experimental data with Protein2PAM model predictions, we generated sequence logos to visualize PAM preferences from HT-PAMDA datasets (Fig. 5c). For each enzyme, we identified all fournucleotide PAM sequences (positions 5–8) that showed cleavage activity (*k >* 5×10^−5^). Each four-nucleotide sequence was weighted by its rate constant, and these weighted sequences were used to calculate a matrix of nucleotide counts, per position. The counts were normalized to frequencies summing to 1.0 per PAM position and then converted to information content for visualization using Logomaker.

## Data and code availability

The training dataset of PAMs was obtained from the CRISPR-Cas Atlas. The Protein2PAM code will be made available upon publication at https://github.com/Profluent-AI/protein2pam, and the machine learning models can be freely accessed through our web server at https://protein2pam.profluent.bio.

## Acknowledgments

We acknowledge funding from the Natural Sciences and Engineering Research Council of Canada (NSERC) Postgraduate Scholarship-Doctoral (PGS D – 567791 to R.A.S.), the Kayden-Lambert MGH Research Scholar Award 2023-2028 (B.P.K.), and National Institutes of Health (NIH) grants DP2CA281401 (B.P.K.), and P01HL142494 (B.P.K.).

## Contributions

S.N., A.B., and A.M. conceived the project. S.N. built the training dataset. A.B. trained the Protein2PAM models. S.N. and A.B. performed the computational experiments. A.N. and S.N. developed the webserver. J.A.R. assisted with structural analysis. G.O.E. and E.H. prepared NmeCas9 variant plasmids for HT-PAMDA with oversight from J.G. N.K. performed HT-PAMDA experiments and R.A.S. assisted N.K. with HT-PAMDA data analysis with oversight from B.P.K. P.C. and A.J.M. provided critical feedback. S.N. prepared the manuscript with input and contributions from A.B. All authors contributed to writing and/or reviewing the final draft of the manuscript.

## Corresponding authors

Correspondence to Stephen Nayfach or Ali Madani.

## Supplementary Information

**Fig. S1.**
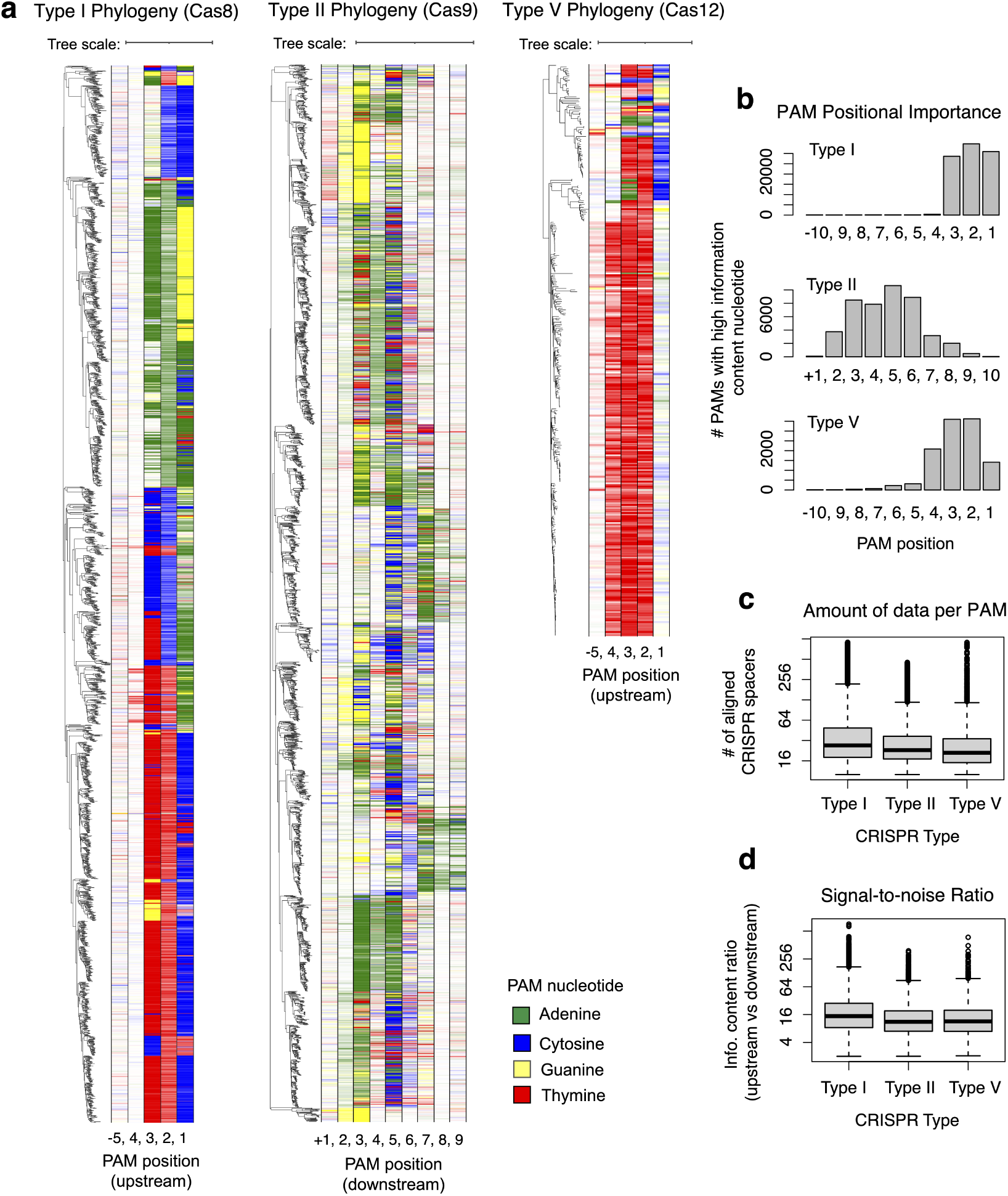
Phylogenetic distribution of PAMs from the CRISPR-Cas Atlas. (a) Phylogenetic trees were built for Cas8, Cas9, and Cas12 proteins. Proteins were first clustered using MMseqs2 (25) at 70% identity for Cas8 and Cas9 and at 95% identity for Cas12. Phylogenetic trees were built using FastTree (26) and visualized using iToL (27). Colored strips indicate the information content at PAM positions. (b) Distribution of high-information content positions across PAMs from Type I, II, and V systems. In Type I systems, the PAM is predominantly restricted to positions −1 to −3 relative to the protospacer, while in Type II systems, the distribution of high information content PAM positions is more variable. (c) Distribution of the number of spacers aligned to virus and plasmid genomes for PAMs predictions from the CRISPR-Cas Atlas. (d) Signal-to-noise ratio comparing nucleotide conservation upstream and downstream of the protospacer for PAMs predictions from the CRISPR-Cas Atlas. In Type II systems, a downstream motif is expected, while in Type I and V systems, the motif is upstream. Bioinformatic PAM predictions are based on a high number of aligned CRISPR spacers, resulting in strong signal-to-noise ratios and providing a robust training dataset for Protein2PAM.

**Fig. S2.**
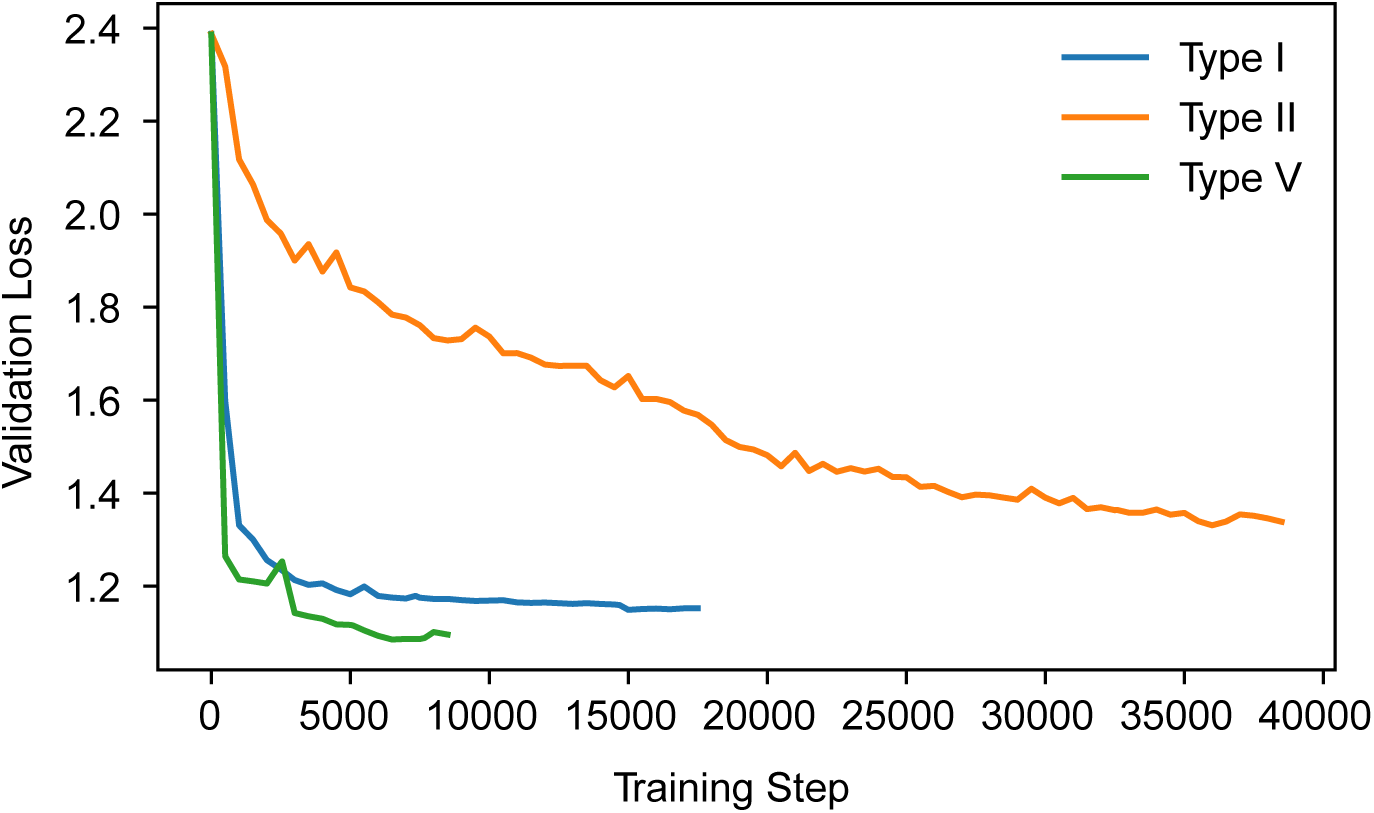
Training loss curves for Protein2PAM models. Each Protein2PAM model predicts nucleotide distributions at 10 PAM positions based on inputted Cas proteins. The architecture integrates a pre-trained 650M-parameter transformer encoder and a 2-layer MLP head. The Type I and V models very quickly converged to their minimum loss, while the Type II model took much longer to optimize. These training dynamics mirror the cross-validation results and show that it is much more challenging to model PAMs for Type II systems than for Type I or Type V systems.

**Fig. S3.**
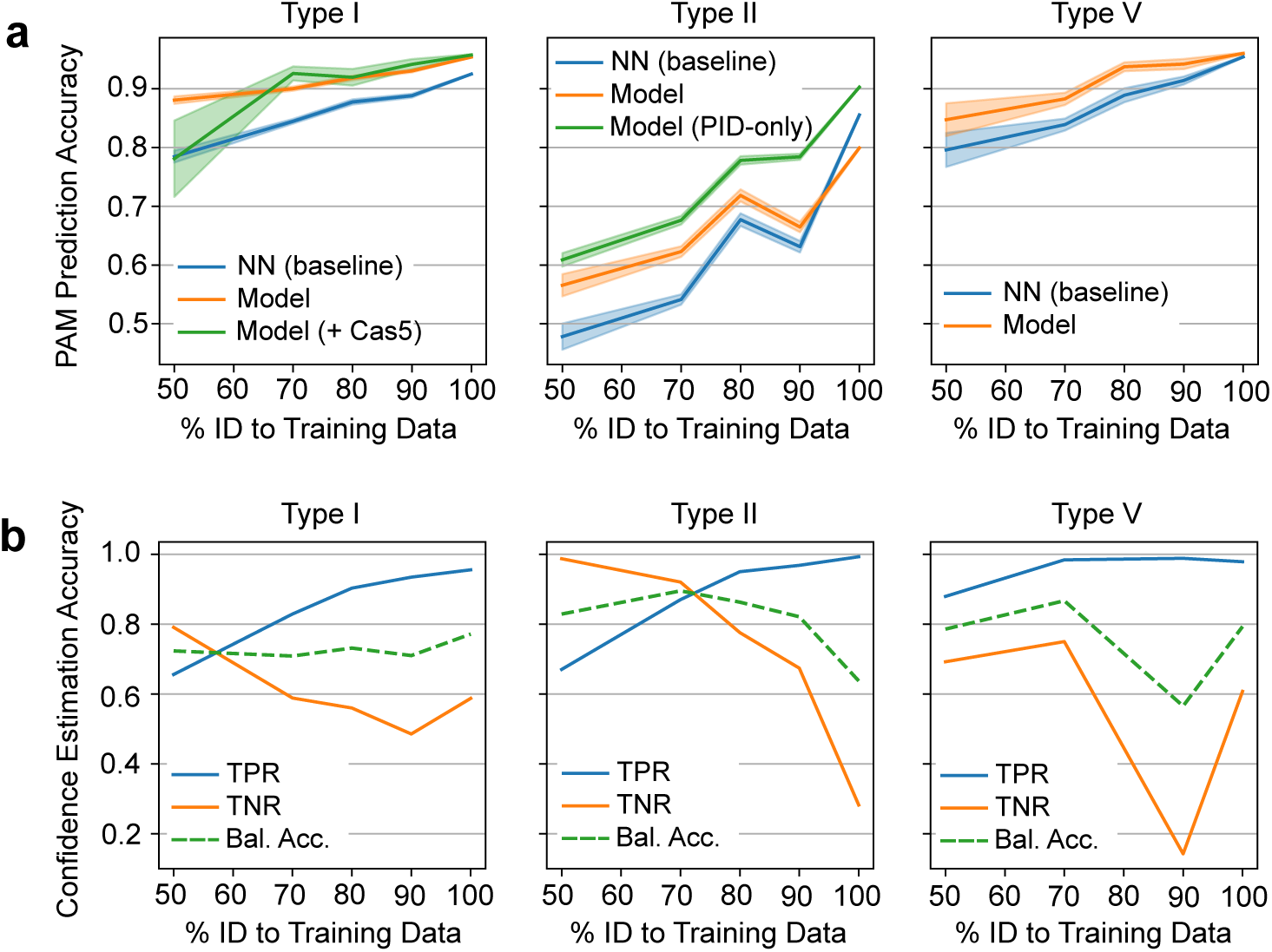
Cross-validation accuracy for Protein2PAM models. (a) PAM prediction accuracy. Each panel indicates the cosine similarity between true and predicted PAMs as a function of distance from the training data. Neural models consistently outperform a baseline in which a sequence is assigned the PAM of the nearest neighbor (NN) in the training dataset. The Cas9 PID-only model outperforms the Cas9 full-sequence model for Type II systems. The Cas8-only model outperforms the Cas8+Cas5 model for Type I systems. (b) Confidence prediction accuracy. True positive rate (TPR), true negative rate (TNR), and balanced accuracy (Bal. Acc.) when determining if a PAM prediction result is high-confidence or not. High-confidence predictions are defined as those with accuracy above 0.80, while accuracy is defined as the cosine similarity between predicted and true PAM. Especially for Type II systems, the confidence model accurately discriminates between accurate and inaccurate Protein2PAM predictions.

**Fig. S4.**
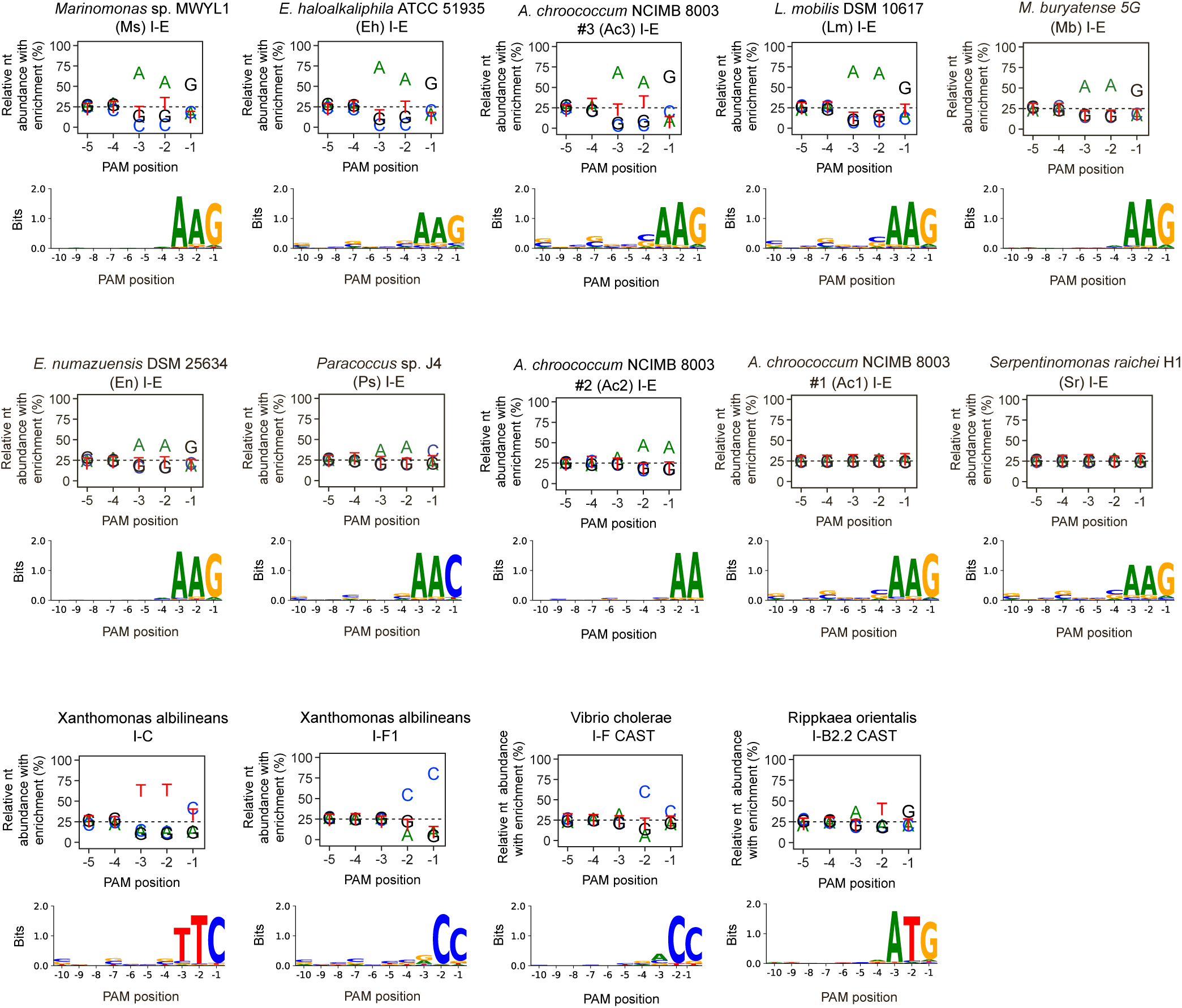
Concordance with experimentally determined PAMs for diverse Type I CRISPR systems. Wimmer et al. characterized PAMs for diverse Type I systems using a rapid cell-free protocol, PAM-DETECT (30). Top panels show nucleotide-enrichment plots from Wimmer et al. for 14 Type I systems subjected to PAM-DETECT. Bottom panels show Protein2PAM predictions for the corresponding Type I systems using Cas8 proteins as input. For two Type I-E systems (Ac1 and Sr), the cell-free assay failed to identify a PAM due to low binding affinity, whereas Protein2PAM was able to confidently predict both PAMs as AAG.

**Fig. S5.**
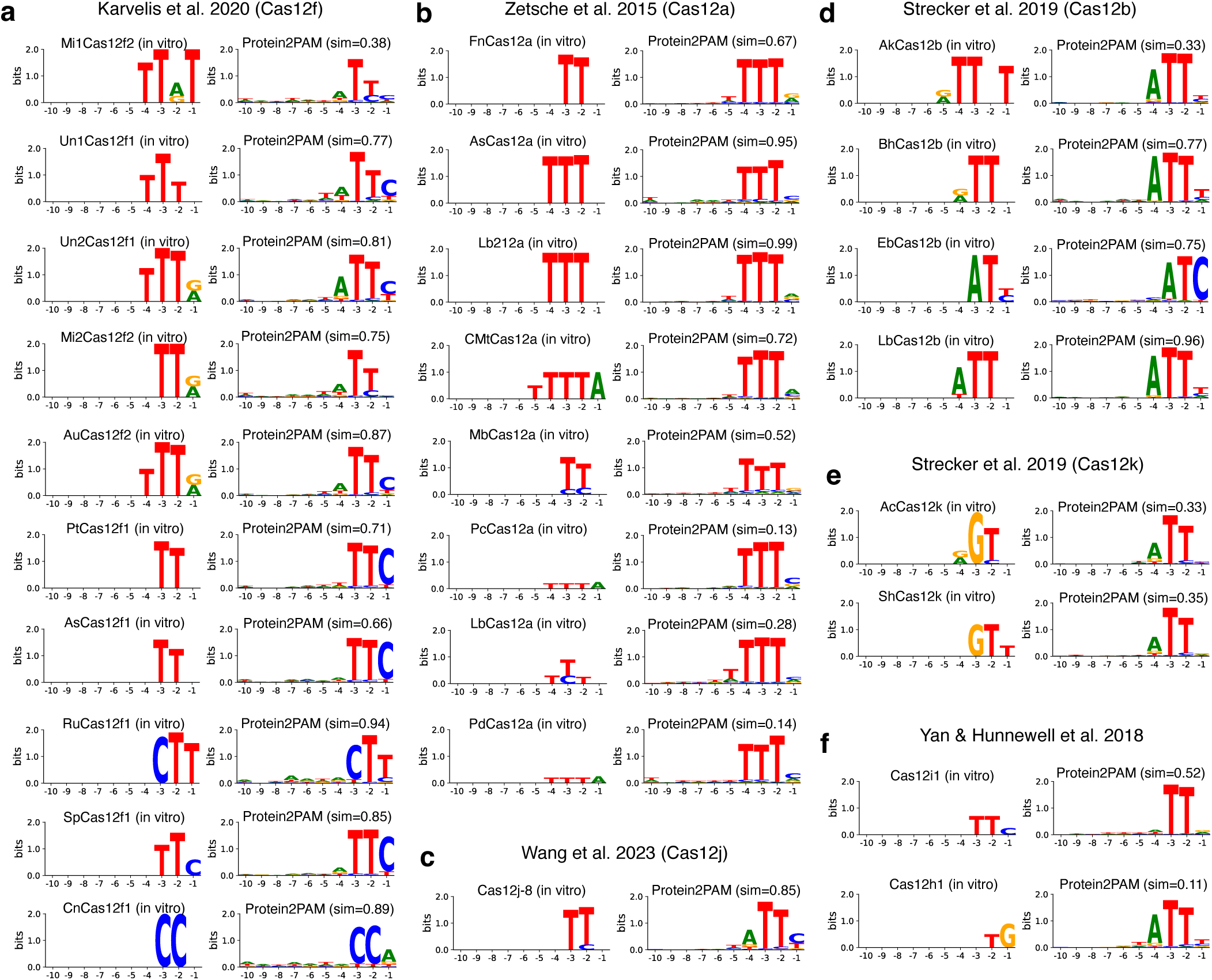
Concordance with experimentally determined PAMs for diverse Type V CRISPR systems. Protein2PAM was applied to 45 Cas12 proteins from 12 different published studies. Protein2PAM predictions were compared to the experimentally determined PAMs using the cosine similarity metric. Figure panels indicate Protein2PAM predictions for 27 of 45 Cas12 proteins. (a) Evaluation of PAM predictions for Cas12f proteins (60). (b) Evaluation of PAM predictions for Cas12a proteins (55). (c) Evaluation of a PAM prediction for Cas12j (61). (d) Evaluation of PAM predictions for Cas12b proteins (57). (e) Evaluation of PAM predictions for Cas12k proteins (62). (f) Evaluation of PAM predictions for Cas12i and Cas12h proteins (4).

**Fig. S6.**
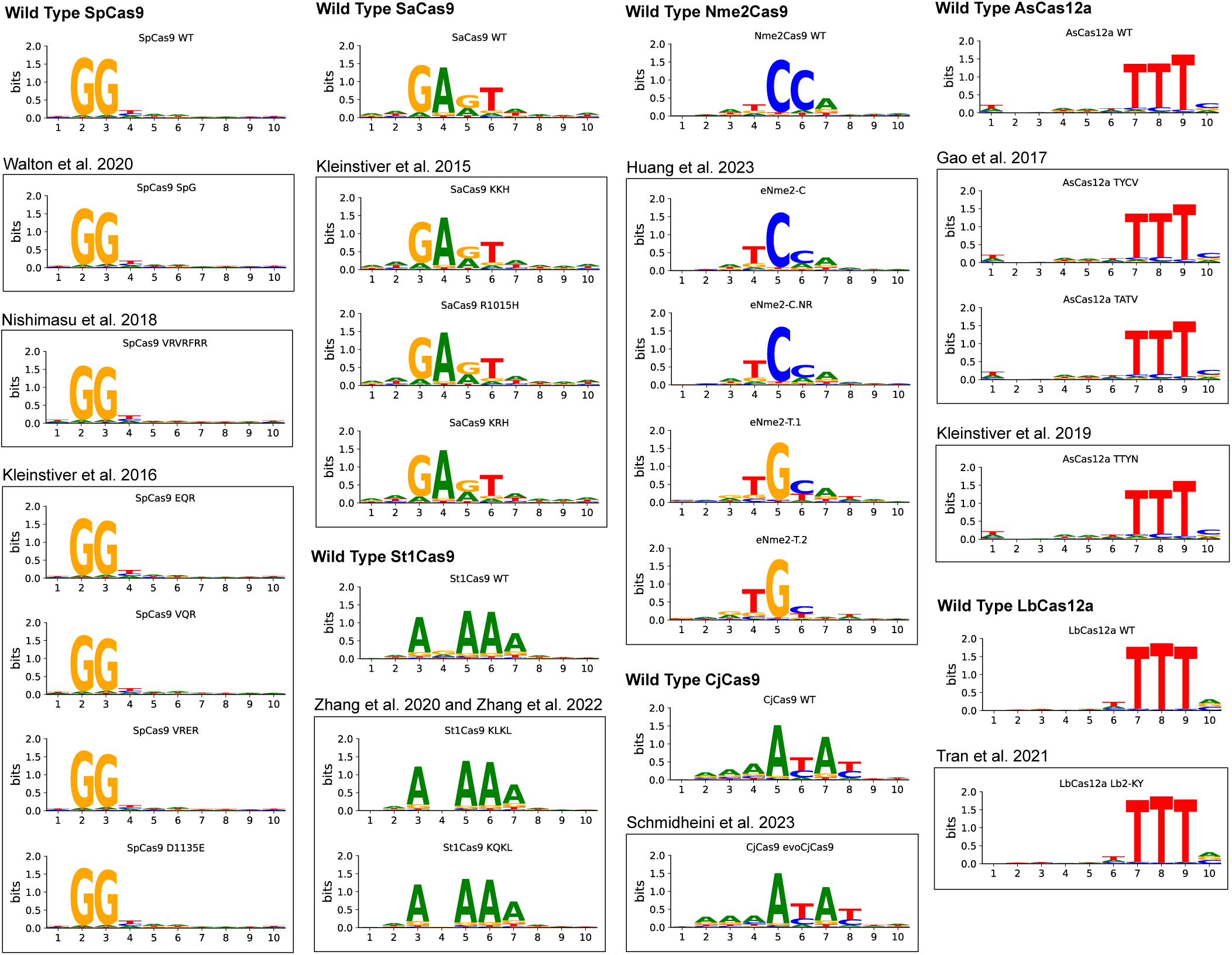
Protein2PAM predictions for previously-engineered Cas enzymes with altered PAMs. We tested Protein2PAM on 20 engineered Cas9 and Cas12 proteins with altered PAM specificities from 10 studies (Methods). These included variants of SpCas9 (7, 8, 11), SaCas9 (10), St1Cas9 (66), Nme2Cas9 (13), CjCas9 (37), and Cas12a (9, 67, 68). In most cases, the model predicted the same PAMs as the wild-type counterparts with the exception of an Nme2Cas9 variant where Protein2PAM correctly predicted a shift from N_4_CC to N_4_CN.

**Fig. S7.**
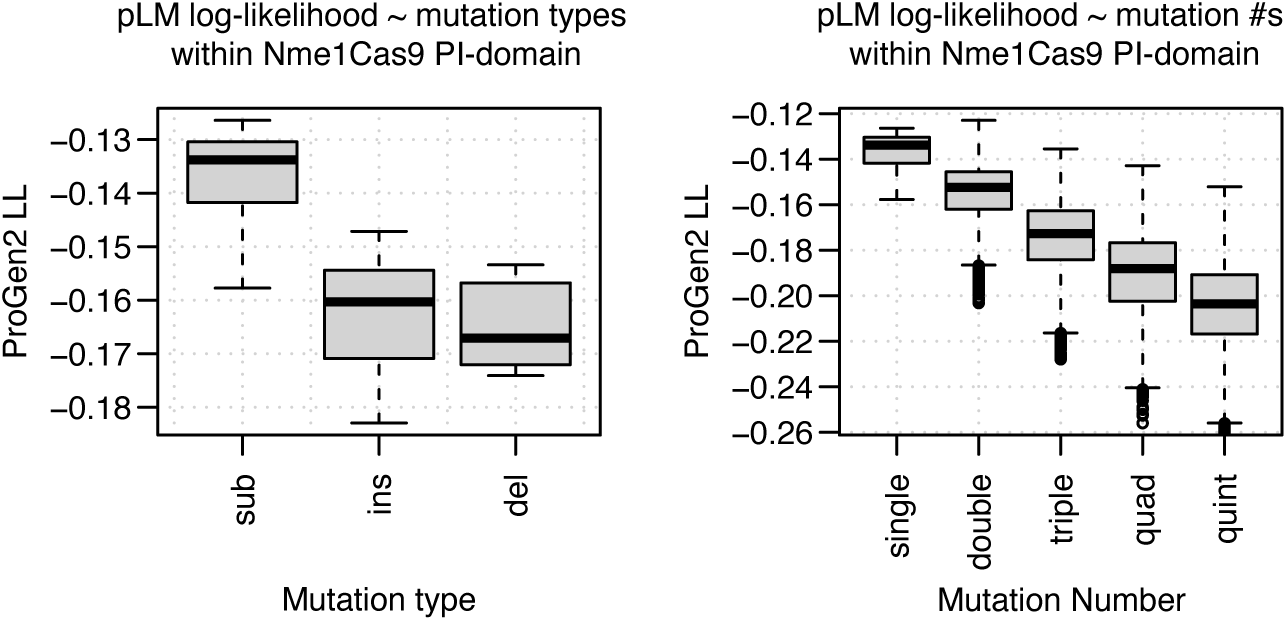
Protein language model scores for Nme1Cas9 mutants. Progen2 fine-tuned on the CRISPR-Cas Atlas was applied to different types of single mutants (left) or mutants with up to five substitutions (right). The language model predicts a decrease in fitness with mutational load and with either insertions or deletions.

**Fig. S8.**
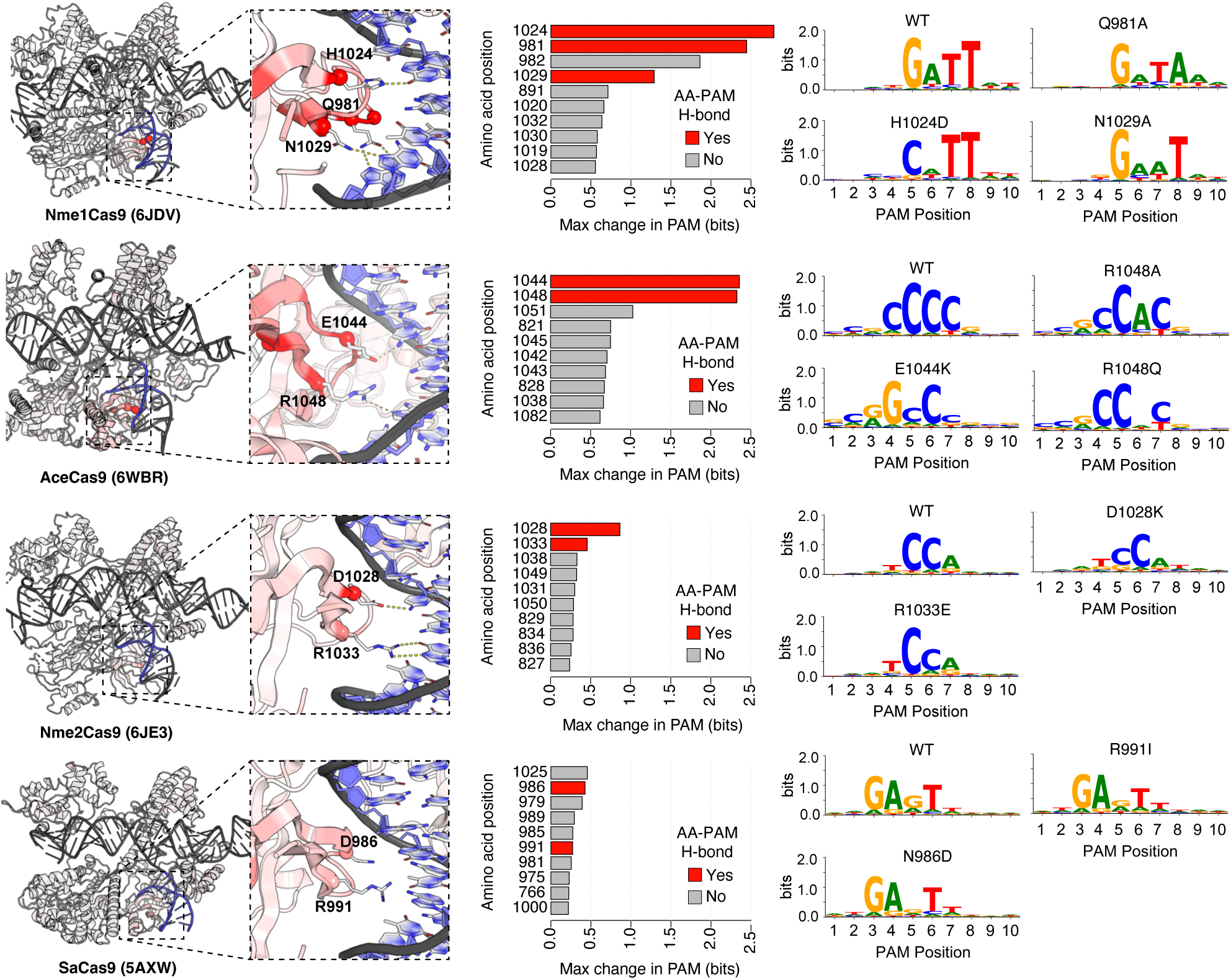
PAM specifying amino acids make hydrogen bonds with specific PAM nucleotides. Four Cas9 crystal structures are shown. Positions are colored by the maximum predicted change to the PAM, in bits, after *in silico* saturation mutagenesis and evaluation with Protein2PAM. Top ranked positions are shown in barplots that result in the greatest predicted change in the PAM after *in silico* saturation mutagenesis. Positions highlighted in red make hydrogen bonds with specific PAM nucleotides in the corresponding crystal structures.

**Fig. S9.**
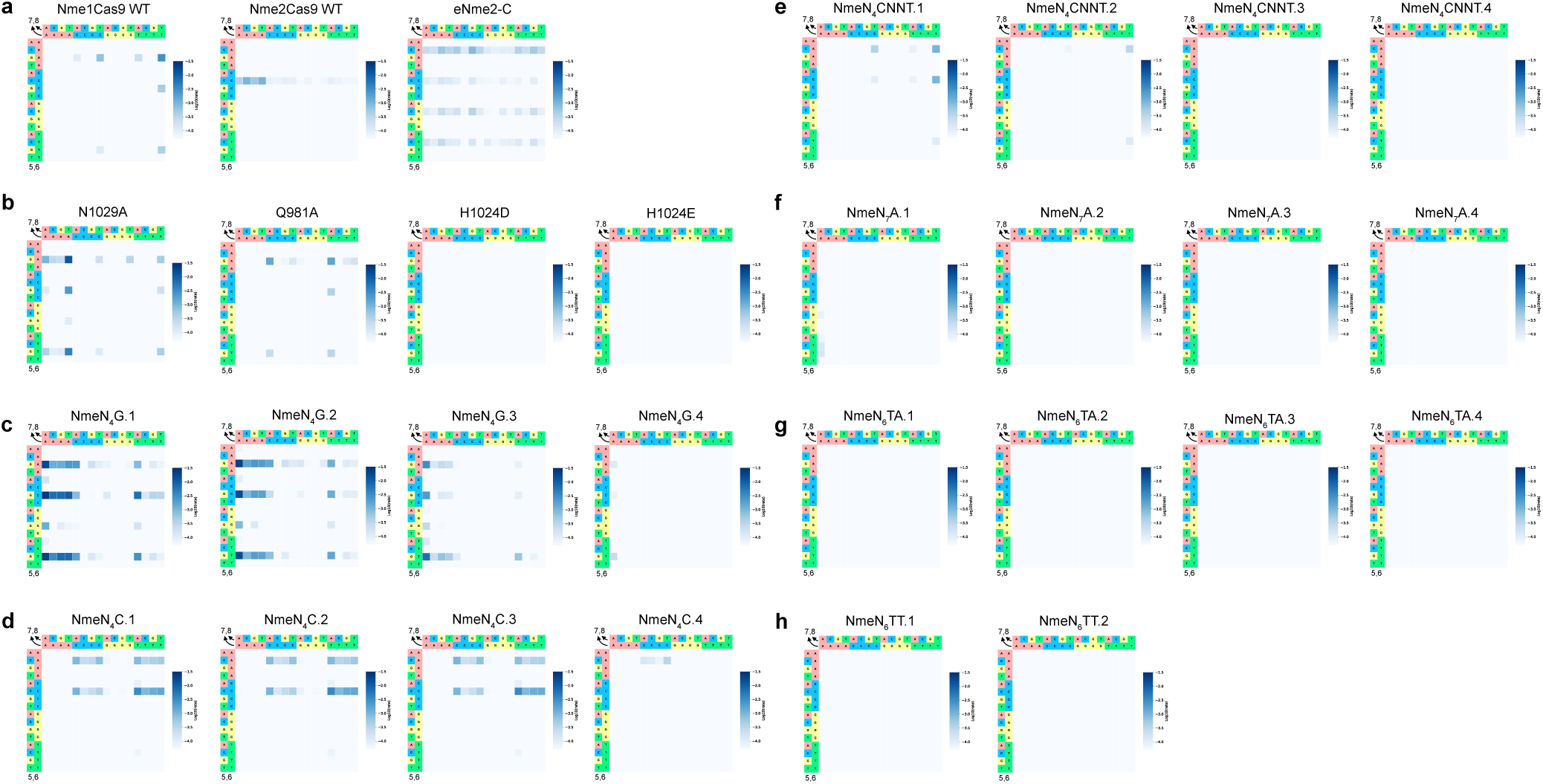
Heatmaps of rate constants from HT-PAMDA data for computationally evolved enzymes. Each panel shows rate constants for Cas9-mediated cleavage at PAM positions 5-8. (a) Wild type and engineered enzymes. (b) Four single amino acid variants. (c-h) 22 Computationally evolved enzyme variants. Overall 11 variants displayed activity in the HT-PAMDA assay with 6 exceeding that of wild-type enzymes. (c) Variants computationally evolved towards N_4_G PAMs. (d) Variants computationally evolved towards N_4_C PAMs. (e) Variants computationally evolved towards N_4_CNNT PAMs. (f) Variants computationally evolved towards N7A PAMs. (g) Variants computationally evolved towards N6TA PAMs. (h) Variants computationally evolved towards N6TT PAMs. See Table S8 for the complete set of HT-PAMDA data for all enzymes.

**Fig. S10.**
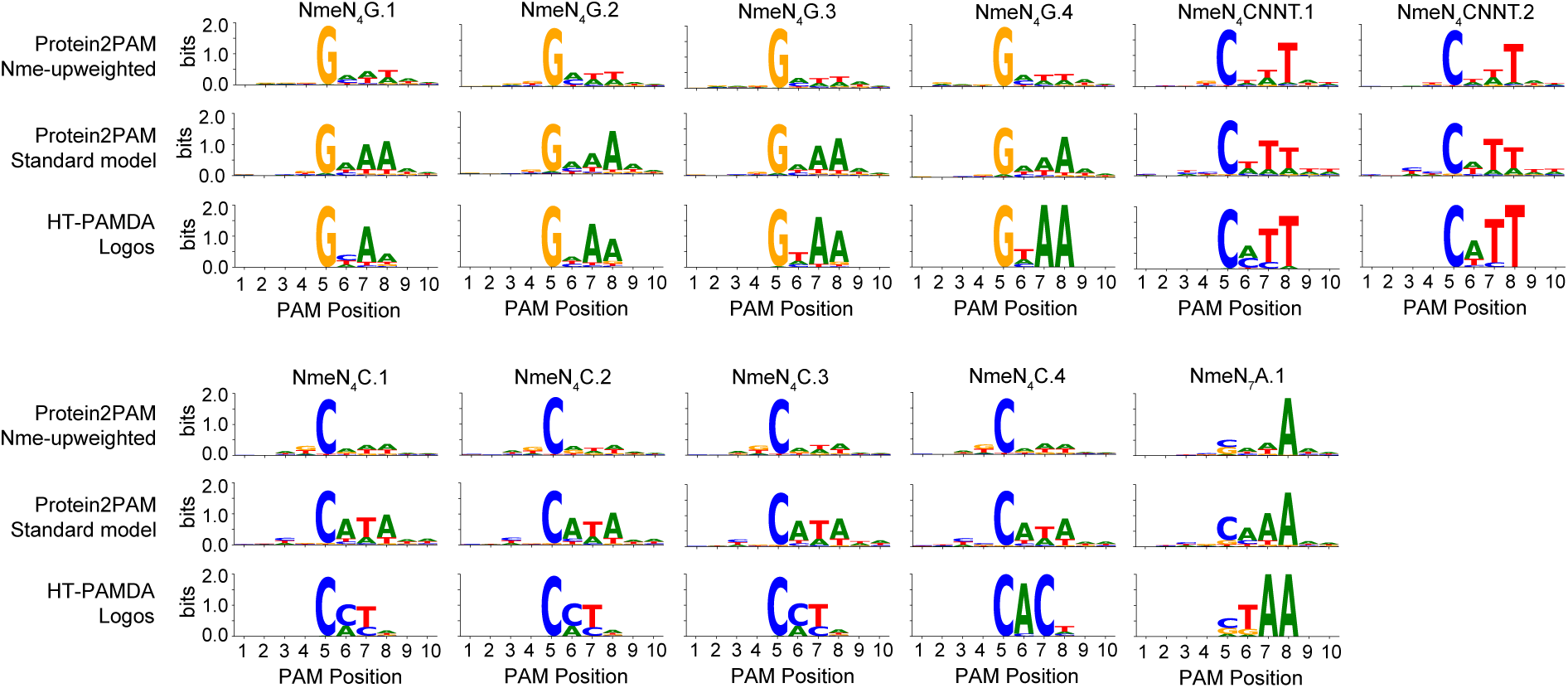
Comparison of PAM predictions between Protein2PAM models for computationally-evolved, active enzymes. Protein names are indicated by plot titles. The first row displays PAMs predicted by the Protein2PAM model used as an oracle during computational evolution with MCMC. In this model, NmeCas9 orthologs were upweighted among the protein:PAM pairs used for training. The second row displays PAMs predicted by the standard Protein2PAM model, which did not include upweighting for NmeCas9 orthologs. Both Protein2PAM models utilized Cas9 PAM-interacting domain sequences for training and inference. The third row displays PAM logos derived from experimental data. The standard Protein2PAM model, which was not used as an oracle in the MCMC process, shows better alignment with the experimental data. The reduced performance of the Nme-specialized model is likely due to extended MCMC trajectories that resulted in overfitting of sequences to the model’s own PAM predictions.

**Fig. S11.**
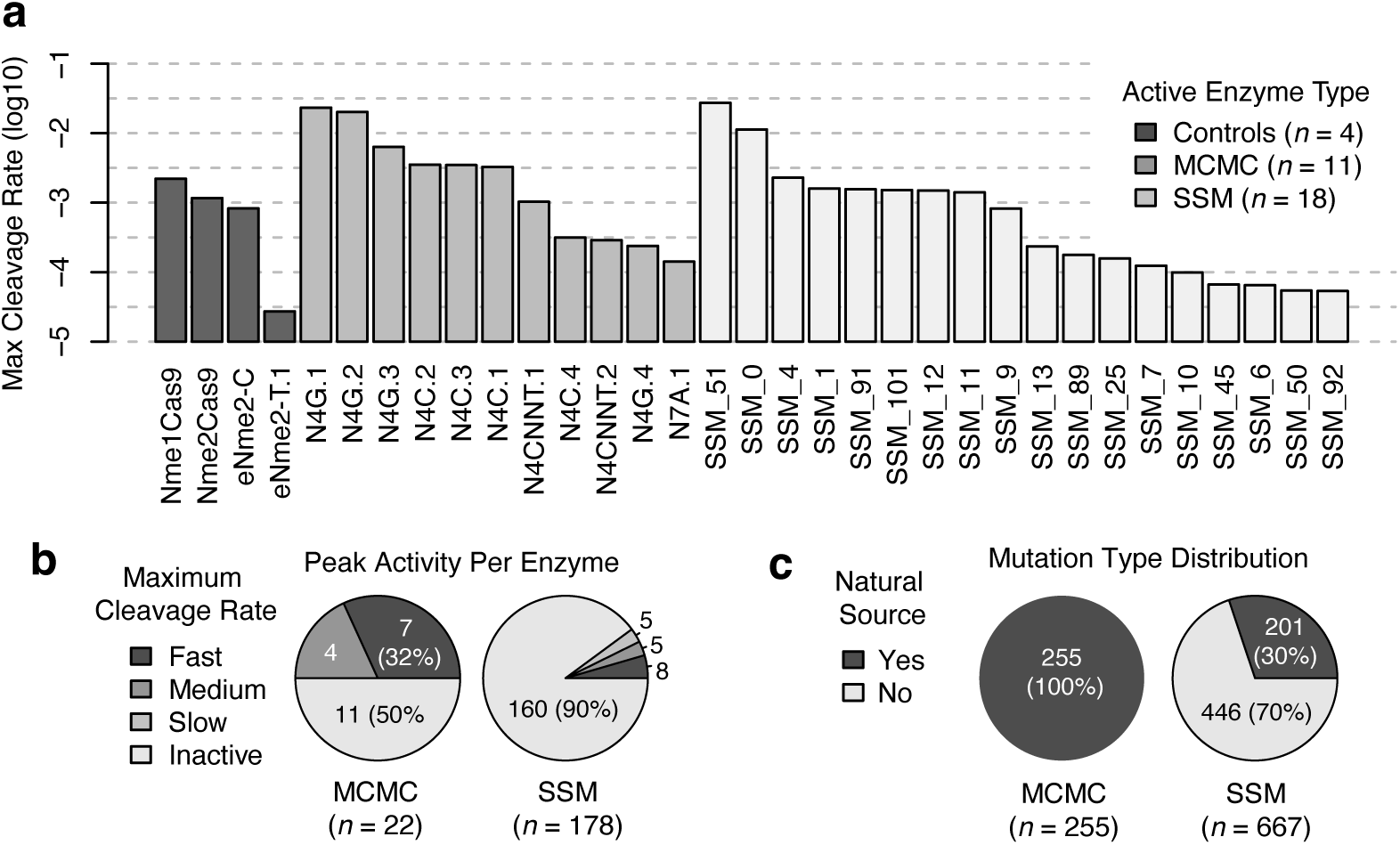
Peak activity of NmeCas9 enzymes across PAMs in HT-PAMDA assay. (a) Each bar represents a single enzyme experimentally characterized by HT-PAMDA. Control enzymes include Nme1Cas9 WT, Nme2Cas9 WT, and eNme2-C and eNme2-T.1 (13). The MCMC sequence category includes 11 active enzyme variants computationally evolved towards six target PAMs. The SSM sequence category includes 18 active enzyme variants containing up to 5 combinations of 102 predicted PAM-specifying mutations. The vertical axis indicates the maximum cleavage rate, *k*, of each enzyme across all 256 PAM in the library, considering only positions 5-8. Enzymes are ranked by sequence type and then by their maximum cleavage rate. (b) Summary of activity rates across all MCMC (*n* = 22) and SSM (*n* = 178) sequences (Fast: *k >* 1 *×* 10*^−^*^3^, Medium: *k >* 1 *×* 10*^−^*^4^, Slow: *k >* 5 *×* 10*^−^*^5^, Inactive: *k ≤* 5 *×* 10*^−^*^5^). (c) Summary of mutation source across mutations found in MCMC (*n* = 255) or SSM (*n* = 667) sequences. A mutation was classified as natural if it was found in a multiple sequence alignment of natural orthologs within 70% amino acid identity of Nme1Cas9.

